# Microneurography as a Minimally Invasive Method to Assess Target Engagement During Neuromodulation

**DOI:** 10.1101/2022.08.19.504592

**Authors:** Nishant Verma, Bruce Knudsen, Aaron Gholston, Aaron Skubal, Stephan Blanz, Megan Settell, Jennifer Frank, James Trevathan, Kip Ludwig

## Abstract

**Objective:** Peripheral neural signals recorded during neuromodulation therapies provide insights into local neural target engagement and serve as a sensitive biomarker of physiological effect. Although these applications make peripheral recordings important for furthering neuromodulation therapies, the invasive nature of conventional nerve cuffs and longitudinal intrafascicular electrodes (LIFEs) limit their clinical utility. Furthermore, cuff electrodes typically record clear asynchronous neural activity in small animal models but not in large animal models. Microneurography, a minimally invasive technique, is already used routinely in humans to record asynchronous neural activity in the periphery. However, the relative performance of microneurography microelectrodes compared to cuff and LIFE electrodes in measuring neural signals relevant to neuromodulation therapies is not well understood.

**Approach:** To address this gap, we recorded cervical vagus nerve (cVN) electrically evoked compound action potentials (ECAPs) and spontaneous activity in a human-scaled large animal model – the pig. Additionally, we recorded sensory evoked activity and both invasively and non-invasively evoked CAPs from the great auricular nerve (GAN). In aggregate, this study assesses the potential of microneurography electrodes to measure neural activity during neuromodulation therapies with statistically powered and pre-registered outcomes (https://osf.io/y9k6j).

**Main results:** The cuff recorded the largest ECAP signal (p<0.01) and had the lowest noise floor amongst the evaluated electrodes. Despite the lower signal to noise ratio, microneurography electrodes were able to detect the threshold for neural activation with similar sensitivity to cuff and LIFE electrodes once a dose-response curve was constructed. Furthermore, the microneurography electrode was the only electrode to record distinct sensory evoked neural activity (p<0.01).

**Significance:** The results show that microneurography electrodes can measure neural signals relevant to neuromodulation therapies. Microneurography could further neuromodulation therapies by providing a real-time biomarker to guide electrode placement and stimulation parameter selection to optimize local neural fiber engagement and study mechanisms of action.

## Introduction

Inconsistent local neural target engagement and poor isolation of on- and off-target neural substrates is critically limiting neuromodulation therapies (Heusser et al., 2016; De Ferrari et al., 2017; Verma et al., 2021a). Direct measurements of neural activity during the acute deployment of neuromodulation therapies could 1) guide electrode design and placement to chronically improve consistency of local target engagement and 2) prove critical for understanding on- and off-target neural substrates and therapeutic mechanisms of action (Verma et al., 2021a).

Although these applications make it pertinent to record neural activity from the peripheral nervous system acutely, the invasiveness of conventional cuff electrodes and longitudinal intrafascicular electrodes (LIFEs) limit their clinical utility. In contrast, clinical microneurography electrodes are minimally invasive, as they can be placed percutaneously into a nerve guided by ultrasound or palpation without a surgical window. Microneurography microelectrodes are often made of tungsten and have a smaller recording tip compared to cuff and LIFE electrodes. They are already used in humans to perform acute recordings for diagnostic and research purposes (Macefield, 2020), but have yet to be evaluated for informing neuromodulation therapies. The relative performance of microneurography electrodes, compared to conventional cuffs and LIFE electrodes, in measuring neural signals relevant to neuromodulation therapies is also not well understood.

To understand the relative performance of microneurography electrodes, compared to conventional cuff and LIFE electrodes, we performed a side-by-side comparison of recording electrodes in the peripheral nervous system of pigs, a large animal model (Fig. 1A). We compared each electrode’s ability to measure both synchronous electrically evoked compound action potentials (ECAPs) (Fig. 1B, top) and asynchronous activity that was either sensory evoked or spontaneously occurring (Fig. 1B, bottom). ECAPs represent the recruitment of neurons around the stimulation electrode and are a direct measurement of local on- and off-target neural engagement. Spontaneously occurring activity is a real-time and sensitive measure of changes induced by neurostimulation, sometimes even before physiological changes become evident (Gonzalez-Gonzalez et al., 2021; Chao et al., 2021). Growing interest in modulating spontaneously occurring neural activity outside the period of stimulation (e.g., burst stimulation for chronic pain therapy) will make spontaneously occurring activity a key biomarker in neuromodulation therapies.

**Figure 1:**
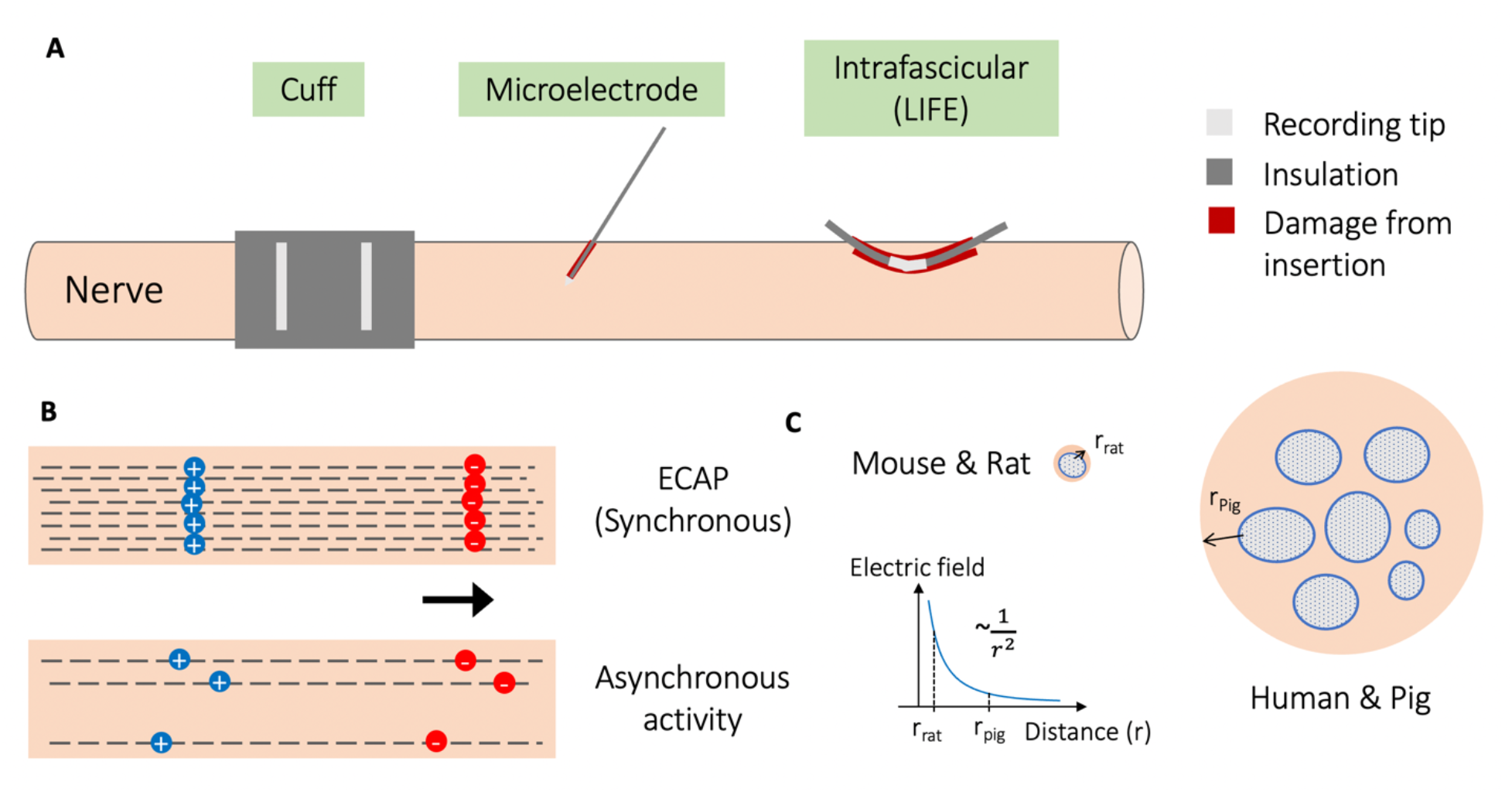
Illustration of three types of recording electrodes, synchronous versus asynchronous neural activity, and effect of electrode-fiber distance on recorded signals. **(A)** Illustration showing the three different types of recording electrodes characterized for their performance in measuring neural signals from the peripheral nervous system. The longitudinal intrafascicular electrode (LIFE) has a larger area of exposed metal for recording than the microneurography microelectrode, and the LIFE causes more damage to tissue upon insertion, indicated by the red blood track. **(B)** Synchronous electrically evoked compound action potential (ECAP) vs. asynchronous neural activity illustrated by number of fibers activated and synchrony of neural activity. Both illustrate a neural signal as a moving dipole with a finite length. This dipole length increases with conduction velocity and fiber size. **(C)** Cartoon illustrating the relevance of scale in interpreting neural recording electrode performance. For example, in the cervical vagus nerve (cVN) for smaller mammals, there is 1) less distance between the neural fiber and the surface of the nerve, where a recording electrode is placed, and hence less falloff in electric field, and 2) there is a thinner layer of perineurium insulation (Pelot et al., 2020), and hence less falloff in electric field. Electric field falloff from the neural source, approximated as a monopole, is proportional to ~1/r^2^, where r is the distance from the neural source to the recording electrode (Plonsey and Barr, 1995).

We performed the experiments in pigs, a human-scaled model, which is critical for the translatability of the findings (Koh et al., 2022). Studies in animal models with smaller peripheral nerves, such as rodents, have shown recordings of spontaneously occurring neural activity made with non-penetrating hook and cuff electrodes (Moncrief and Kaufman, 2006; Silverman et al., 2018; Stumpp et al., 2020). However, this might not be possible in human-scaled nerves (Koh et al., 2022). The cervical vagus nerves (cVNs) in mice and rats are 200-300 *µ*m in diameter with a single fascicle while the cVNs in humans and pigs are 2-3 mm in diameter with many fascicles (Settell et al., 2020; Pelot et al., 2020). The larger diameter of the nerve and thicker low electrical conductivity perineurium and epineurium increase the distance between the firing neuron and the recording electrode (electrode-fiber distance), thereby reducing the recorded signal amplitude (Yoo et al., 2013) (Fig. 1C). As an example, Yoo and colleagues (2013) reported that they had to de-sheath the epineurium of cVN in dogs, thereby reducing electrode-fiber distance, to clearly record C-fiber ECAP components. The larger electrode-fiber distances of nerves in human-scaled models might make it challenging to record asynchronous neural activity with non-penetrating electrodes.

Here, we evaluated microneurography electrodes against cuff and LIFE electrodes in their ability to measure invasively and non-invasively evoked CAPs, sensory evoked neural activity, and spontaneously occurring activity in the peripheral nerves of swine, a large animal model representative of the human peripheral nervous system (Settell et al., 2020). The cVN, a major autonomic nerve of great interest as a neuromodulation therapy target (Johnson and Wilson, 2018), was used as the first model nerve. In addition, the great auricular nerve (GAN), a sensory nerve innervating the ear and implicated in auricular vagus nerve stimulation (aVNS) (Verma at el., 2021a), was used as the second model nerve to evaluate sensory evoked potential and non-invasively evoked CAP recordings. We examined the effects of reference electrode placement on neural signal fidelity and artifacts. We noted common non-neural artifacts in electrophysiology recordings, and we discuss appropriate controls to verify neural recording authenticity. Based on the findings, we summarize considerations and recommend minimally invasive microneurography as a technique that can be used acutely during the deployment of neuromodulation therapies to inform electrode placement, electrode design, and selection of stimulation parameters to optimize local neural target engagement. Understanding on- and off-target neural fibers engaged is a critical step in investigating therapeutic mechanisms of action and could deepen our understanding of both peripheral and central mechanisms of neuromodulation therapies.

## Methods

In this study, we characterized the ability of three electrodes to measure neural activity in the peripheral nervous system. We characterized 1) cuff electrodes, 2) LIFE electrodes, an intrafascicular electrode used in pre-clinical studies (Yoshida and Stein, 1999; Nicolai et al., 2020), and 3) microneurography electrodes, a clinical microelectrode used routinely in humans (Macefield, 2021). The cVN, a major autonomic nerve innervating organs in the thorax and abdomen as well as muscles in the throat, and the great auricular nerve (GAN), a sensory nerve innervating the ear and periauricular region, were used as model nerves due to their relevance in existing peripheral neuromodulation therapies. The cVN is targeted during invasive vagus nerve stimulation (VNS) for epilepsy and depression (Johnson and Wilson, 2018) and the GAN is involved in non-invasive aVNS (Cakmak et al., 2017; Verma et al., 2021a).

### Pre-registration

Pilot experiments (n=10) were conducted to refine the surgery and establish the study protocol. The study was then pre-registered (https://osf.io/y9k6j) with sequentially defined primary, secondary, and exploratory outcomes, statistical comparisons, analysis parameters, and sample size (n=6). The confirmatory experiments were conducted after pre-registration with no exclusion of subjects.

### Recording and stimulation electrodes

Photographs of the recording electrodes are in Supplementary Material 1.

The microneurography electrodes were fabricated of tungsten with epoxylite insulation. They had an impedance of 0.8-1.2 M*Ω* at 1 kHz, shaft diameter of 250 *µ*m, and rounded tip diameter of ~5 *µ*m (part #UNA40GCT, FHC Inc., ME, USA).

The GAN recording cuffs were identical to the GAN stimulation cuffs and were fabricated in-house with two platinum wires 127 *µ*m (0.005”) in diameter, separated by ~1 mm, glued (using a silicone-based glue) into a split silicone tube of 0.75 mm inner diameter with a total length of 3 mm (Fig. 2B).

**Figure 2:**
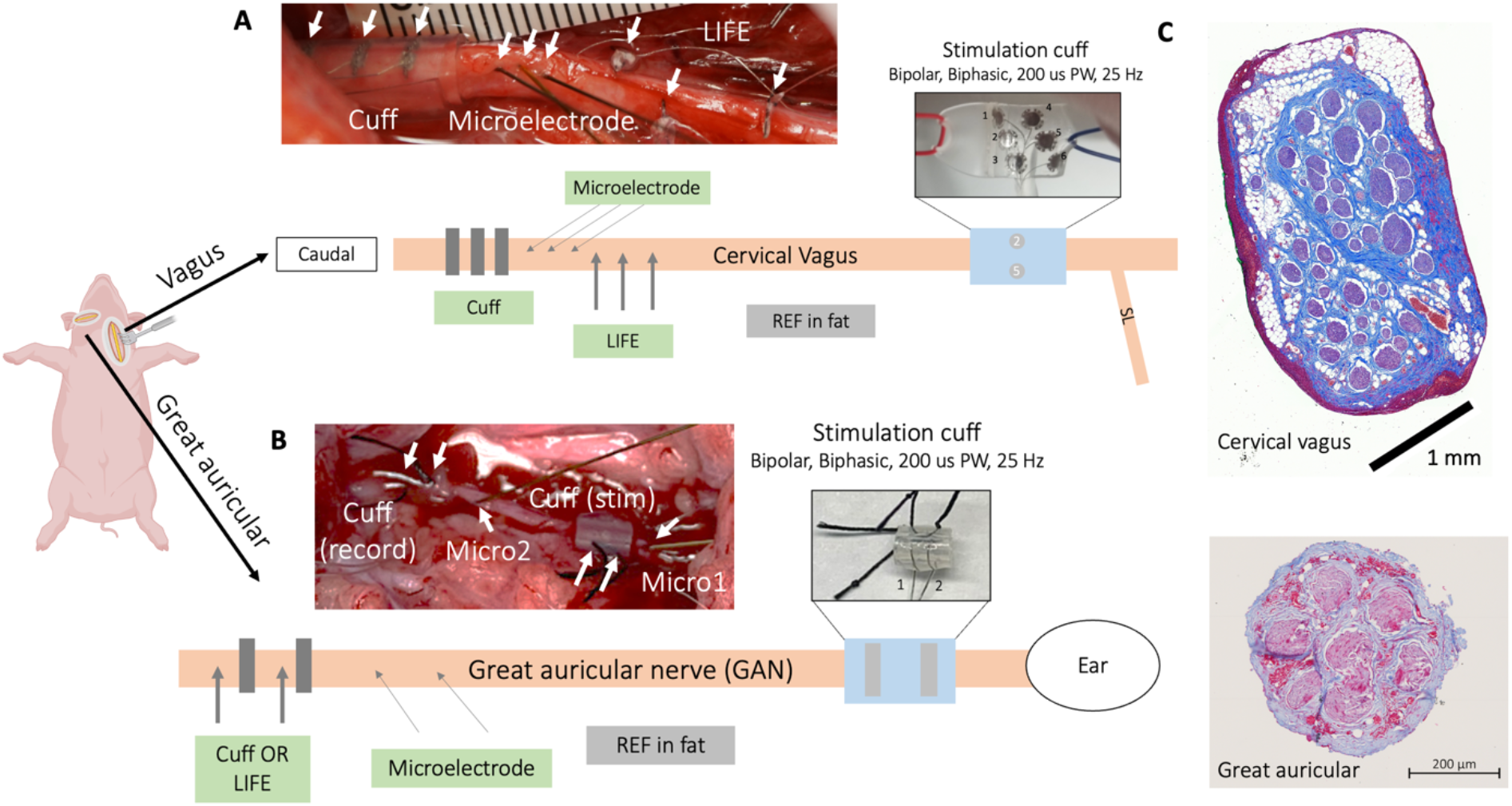
Experimental setup and surgical preparation. **(A)** Cervical vagus nerve (cVN) instrumented with bipolar stimulation cuff (contact 2 and 5) and, at least 4 cm caudal, three replicates each of three different recording electrodes: three longitudinal intrafascicular electrodes (LIFEs), three microneurography electrodes, and a three-contact cuff. All recording electrodes were referenced to fat (off-nerve local tissue reference). **(B)** Great auricular nerve (GAN) instrumented with bipolar stimulation cuff and two replicates each of two different recording electrodes: two microneurography electrodes, and either a two-contact cuff or two LIFE electrodes (alternated by subject). All recording electrodes were referenced to fat (off-nerve local tissue reference). **(C)** Histology of the cVN and the GAN under the stimulation cuff illustrating the relative difference in size and fascicular organization. The cVN is approximately 5x larger than the GAN and contains a greater number of fascicles. GAN histology from additional subjects is shown in Supplementary Material 2.

The cVN recording cuff was fabricated of platinum and silicone with three cylindrical contacts 1 mm in width, 3 mm apart edge-to-edge, and 2 mm from the edge of the silicone. Each of the three cylindrical contacts was split into seven contacts (1 × 1 mm^2^ size), 0.5 mm apart edge-to-edge, with all seven contacts electrically connected to improve mechanical flexibility. The cVN recording cuff had an inner diameter of 3 mm (Ardiem Medical Inc., PA, USA).

The cVN stimulation cuff was fabricated of platinum-iridium and silicone. It was a six-contact cuff with each contact 2 mm in diameter (Blanz et al., 2022). Contacts 2 and 5 were used as the bipolar pair for stimulation (Fig. 2A).

The LIFE electrodes were fabricated in-house of platinum-iridium wire, 102 *µ*m (0.004”) diameter with insulation, 51 *µ*m (0.002”) without insulation, and an exposed window length of ~2 mm (Nicolai et al., 2020; Blanz et al., 2022). A curved insertion needle was used to ‘sew’ the LIFE electrode into the nerve. The impedance of the LIFE electrodes was 1-10 k*Ω* at 1 kHz.

### Acute swine experiments

We used n=6 (3M/3F) domestic swine, 50-60 kg, ~3 months old in the confirmatory experiments. All animal care and procedures were approved by the University of Wisconsin-Madison Institutional Animal Care and Use Committee (IACUC).

#### Anesthesia

An intramuscular injection of Telazol (6 mg/kg) and xylazine (2 mg/kg) was used to induce anesthesia. The animals were intubated and mechanically ventilated, and the surgical anesthetic plane was maintained with inhaled isoflurane (1-2%) and intravenous fentanyl (12-30 *µ*g/kg/hr), administered with lactated Ringer’s solution (LRS), for analgesia.

Several hours into the experiment, an onset of tremors was observed in the animals. Pigs are known to be especially prone to tremor during anesthesia (Ringer et al., 2016). The tremors were observed despite confirmation of the surgical anesthetic plane by means of nasal septum pinch, jaw slackness, unresponsiveness to corneal reflex, and verification of other physiologic parameters such as temperature, blood glucose, and blood pH. The tremors persisted despite increased dosing of isoflurane and fentanyl. We found either intramuscular injection of Telazol (4-6 mg/kg) or intravenous ketamine (10 mg/kg/hr) eliminated the tremors. The effects of Telazol lasted for ~2 hours while ketamine effects on tremor washed in and out in ~15 minutes.

A muscle paralytic, vecuronium, was administered intravenously during cVN recordings to avoid EMG artifacts in the neural recordings from electrically evoked muscle movement (Yoo et al., 2013; Nicolai et al., 2020; Blanz et al., 2022) or ongoing tremors – without cardiac blunting effects. We delivered vecuronium as a 0.1 mg/kg bolus over 1 minute followed by 1-1.5 mg/kg/hr constant rate infusion.

#### Physiological recordings and data analysis

We recorded heart rate from a pulse transducer (AD Instruments, Sydney, Australia) and EKG (AD Instruments, Sydney, Australia), blood pressure from a femoral artery catheter (Millar, Houston, TX, USA), temperature from a rectal probe (Physitemp, Clifton, NJ, USA), and SpO2 from a pulse oximeter (Nonin, Plymouth, MN, USA) applied on the tongue. The signals were recorded with a PowerLabs 8/35 (AD Instruments, Sydney, Australia). Stimulation timings were synced with the physiological recordings, using a TTL signal line from the stimulator to the PowerLabs, for further analysis of stimulation-evoked physiological changes.

Stimulation-evoked heart rate changes were calculated as the difference between the mean heart rate 1-3 s before stimulation and the maximum change in heart rate during stimulation (Blanz et al., 2022).

#### Cervical vagus nerve preparation and electrophysiology

ECAPs from invasive stimulation and spontaneous activity were measured from the cVN (Fig. 2A, C). With the subject in supine position, a midline approach was used to access the left carotid sheath. The carotid artery was mobilized and carefully retracted to minimize obstruction to blood flow. The cVN was exposed for a length of 9-12 cm and instrumented with a bipolar stimulation electrode caudal to the superior laryngeal branching off the nodose ganglion (Settell et al., 2020). Three LIFE electrodes, three microneurography electrodes, and a cuff electrode with three recording contacts were instrumented on the nerve caudal to the stimulation electrode. A separation of >4 cm was kept between the stimulation electrode and the closest recording electrode. The order of the recording electrodes (cuff, LIFE, and microneurography) were randomized, without replacement, per subject to reduce bias arising from distance between the stimulation and recording electrode, which is known to affect the ECAP (Parker et al., 2020). A reference electrode, similar in design to the recording LIFE electrodes, inserted in superficial fat was used to reference the cuff and LIFE electrodes while a microneurography electrode inserted in superficial fat was used to reference the microneurography electrodes. Superficial fat was selected for its similarity in composition to the fatty tissue of nerves, thereby best matching the mechanical and electrochemical environment of the recording electrodes (Ludwig et al., 2008). An off-nerve reference also allowed for post-hoc virtual re-referencing between all recording electrode pairs in each reference group. The reference electrode site was selected at a distance approximately equidistant from the stimulation electrode to the recording electrodes to best match, and ‘subtract’, the representation of the stimulation artifact. In addition, several recordings were collected with on-nerve ‘bipolar’ and ‘tripolar’ referencing to compare the three referencing methods.

A Tucker-Davis Technologies (TDT) electrophysiology system was used for stimulation and recording. The analog front-end and digitizer were on the battery-powered Subject Interface Module (SIM, TDT Inc., FL, USA). Data were digitized at 25 kHz with a 28-bit Sigma-Delta analog to digital converter (ADC) (45%/11.25 kHz low-pass anti-aliasing filter) without amplification. High impedance (microneurography electrodes) and low impedance (LIFEs and cuff) were recorded into two separate recording cards with an active (ZC32, TDT Inc., FL, USA) and passive (S-Box, TDT Inc., FL, USA) head stage, respectively. Stimulation was delivered in bipolar mode with two floating current sources.

ECAPs were recorded during delivery of 750 symmetric biphasic stimulation pulses with a 200 *µ*s pulse width at 25 Hz. These stimulation pulse trains, 30 seconds each, with amplitudes between 0 and 10 mA were presented in a randomized order with 60 seconds between subsequent trains. Spontaneous activity recordings were made before electrical stimulation was delivered.

#### Great auricular nerve preparation and electrophysiology

The GAN innervates the auricle and is implicated as an off-target nerve in some aVNS therapies. The GAN is also purported to be part of the therapeutic effect pathway for auricular stimulation-based therapies to treat Parkinson’s disease (Cakmak et al., 2017) and pain (Kaniusas et al., 2019). Therefore, measuring neural activity directly from the GAN could be used to assess both on- and off-target neural engagement during auricular stimulation therapies to inform stimulation electrode location and parameters. The GAN (Fig. 2C, bottom) is morphologically distinct from the cVN (Fig. 2C, top), which is larger and contains a greater number of fascicles – influencing the spread of electric fields generated by firing neurons and consequent neural recordings. Characterizing recording electrode performance in both these nerves improves the applicability of our findings. The GAN was also selected because it was more surgically accessible than the primary on-target nerve for aVNS therapies, the auricular branch of the vagus (ABVN), and potentially amenable to percutaneous microneurography under ultrasound, which may be problematic for the ABVN due to its anatomical course.

ECAPs from invasive stimulation and non-invasive stimulation and sensory evoked neural activity in response to sensory stroking were measured from the GAN. The GAN was accessed by the approach described in the following paragraph, which to our knowledge, offers the first surgical description of a procedure specifically accessing and identifying the GAN in swine.

We created a larger than necessary surgical pocket, as our approach was to find the pathway of the sensory GAN and identify the association it had with the auricular motor components of the facial nerve (facial auricular nerves). In the surgical pocket (photograph in Supplementary Material 3), we located and traced either of two landmarks to locate the GAN. We followed the facial auricular nerves dorsally in search of the GAN traveling in association. Alternatively, we followed the lateral auricular vein ventrally, to where the nerves (sensory and motor) and vein course together. The skin and subcutaneous fat were incised from the medial posterior margin of the ramus. A small notch was palpated at this point and indicated the approximate level of the stylomastoid foramen. Here, the facial nerve exited and divided into its various branches (Getty, 1975) (more details on facial nerve anatomy are provided in the Supplementary Material 4). The incision continued dorsally following a line posterior to the temporomandibular articulation and up to the medial base of the ear just inferior to the medial crus of the helix. This allowed us to follow the lateral vein through the base of the ear as a landmark for finding the sensory input for the exterior skin of the auricle (the GAN). This incision exposed the superficial musculoaponeurotic system (SMAS), a superficial fibromuscular layer that integrated with the superficial temporal fascia dorsally and platysma ventrally. The SMAS layer was divided along the posterior margin of the ramus and posterior margin of the temporomandibular articulation/zygomatic process to expose the subfascial level adipose tissue. This underlying adipose tissue along the posterior margin of the temporomandibular articulation was the location where we were able to isolate the GAN (more details on GAN anatomy are provided in the Supplementary Material 5) and branches of the facial nerve (Getty, 1975; Lefkowitz et al., 2013; Yang et al., 2015; Sharma et al., 2017). Running in conjunction with these nerves were the lateral/caudal auricular artery and vein, matching the description of Duisit et al. (2017).

To determine which of the exposed nerves were motor or sensory in origin we electrically stimulated each of the nerve branches to evoke a motor response – indicative of a motor pathway (200 *µ*s pulse width symmetric biphasic pulses, 0.02-1 mA, 1 Hz). This is similar to the procedure used to localize a microneurography electrode into the desired fascicle (Macefield, 2021). When the sensory branch of the posterior auricular nerve was identified by no motor response, it was instrumented with a stimulation cuff (Fig. 2B) and recording electrodes (two microneurography electrodes and a two-contact cuff or two LIFE electrodes). The total length of the exposed GAN was ~5 cm. The identification of the sensory GAN was verified by recording sensory evoked responses during stroking of the skin at the base of the auricle. Reference electrodes were placed similarly as described above in the cVN preparation and the same electrophysiology recording setup was used.

Sensory evoked neural activity was recorded by lightly stroking the region, close to the base of the ear, innervated by the GAN (‘on-target’) with a toothbrush (Fig. 7A, green). Each trial lasted 90 seconds: 30 seconds of no stroking, 30 seconds of ‘on-target’ stroking, and 30 seconds of ‘off-target’ stroking (closer to tip of ear, Fig. 7A red) where the GAN is not expected to innervate. The ‘off-target’ stroking served as a control to ensure that the light stroking applied to the ear did not transfer to the recording electrodes as motion artifacts.

Transcutaneous electrical nerve stimulation (TENS) electrodes cut to 2 × 2 cm^2^ in size were applied to the ‘on-target’ region and at the base of the ear through which the main trunk of the GAN courses. Non-invasively evoked CAPs were recorded during application of stimulation waveforms with parameters identical to those used to stimulate the cVN.

Invasively evoked CAPs were recorded during application of stimulation waveforms with parameters identical to those used to stimulate the cVN, except with stimulation amplitudes between 0 and 3-10 mA (determined based on each subject’s motor response threshold). Sensory evoked recordings were performed first, followed by invasively evoked CAP recordings, and lastly non-invasively evoked CAP recordings.

### Electrophysiology data analysis

A custom-built and publicly available Python package, PyeCAP (https://github.com/ludwig-lab/pyeCAP), was used for offline analysis of electrophysiology and physiology data.

#### Filtering

ECAP electrophysiology data was filtered with a high pass 1^st^ order Butterworth filter with a corner frequency at 100 Hz and a low pass Gaussian filter with a corner frequency at 3 kHz. The low pass gaussian filter negates the possibility of introducing filter ‘ringing’ artifact, which may be introduced by a Butterworth filter that has an ‘overshoot’ to an impulse response (Bovik and Acton, 2005). The stimulation artifact approximates an impulse, and an overshoot following a stimulation artifact – caused by inappropriate filter selection – could be mistaken for an ECAP. An additional 60 Hz band stop finite impulse response (FIR) filter constructed with a Hamming window was used on the sensory evoked and spontaneously occurring neural activity electrophysiology data. All filtering was performed on the time series data in both the forward and backward direction to eliminate group delays caused by filtering (Scipy, 2022).

#### Non-functional recording electrodes

As per the pre-registration (https://osf.io/y9k6j), non-functional recording electrodes were identified by a high noise floor, high 60 Hz noise, or a flat ECAP dose-response behavior in comparison to other contacts of the same electrode type. Electrodes deemed non-functional by these criteria were noted (Supplementary Material 6) and removed from analysis.

#### Spike detection

Spike detection for spontaneously occurring activity and sensory evoked recordings was done using a voltage thresholding method. The threshold voltage was set per electrode at six times the standard deviation (SD) from time=1-6 seconds of the time series trace recorded by that electrode added to the mean calculated from the same time window (time=1-6 seconds) to account for baseline offsets. No sensory evoked activity was initiated during time=1-6 seconds, although spontaneously occurring activity was not accounted for. This may have inflated the threshold for spike detection slightly.

#### Detecting authentic ECAPs

ECAPs were plotted by averaging (point-by-point median) the stimulation evoked response across 750 pulses in a particular stimulation train. Averaging multiple stimulation evoked responses reduces uncorrelated noise by a factor of 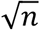, where n is the number of traces averaged (Metcalfe et al., 2018). The averaged ECAP traces were split into time windows by neural fiber type according to the distances between recording and stimulation electrode and published conduction velocities (Erlanger and Gasser, 1937; Manzano et al., 2008).

ECAP authenticity was confirmed by several methods. First, signal propagation delay across recording electrodes (Gaunt et al., 2009) was verified to be in the expected nerve fiber conduction velocity range (Erlanger and Gasser, 1937; Manzano et al., 2008). An artifact (e.g., EMG, motion) could occur at a similar time point as an ECAP but would not show up with a signal propagation delay across spatially separated recording electrodes (Gaunt et al., 2009). Second, a muscle paralytic, vecuronium, was used to abate EMG artifact, which could be mistaken for an ECAP, in the cVN ECAP recordings. Third, transection of the nerve caudal and cranial to the recording electrodes and subsequent recordings were used to confirm the disappearance of the ECAP signal, which is no longer able to propagate along the transected neural pathway from stimulation to recording electrode (Nicolai et al., 2020).

#### Detrending neural signal from stimulation artifact

A small subset of ECAP recordings were contaminated with stimulation artifacts; the neural signal of interest was riding ‘on top’ of a stimulation artifact decay. The ECAP signal had to be separated from the baseline offset of the stimulation artifact to quantify its RMS magnitude accurately. Neural signal was detrended from stimulation artifact using an exponential based ‘fit and subtract’ approach (Harding, 1991; Chakravarthy et al., 2022; Drebitz et al., 2020). A simple method was used as only ECAPs from Subject 6 microneurography electrode recordings during the cVN dose-response experiment were contaminated by stimulation artifacts. A single exponential decay of the form:

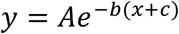

with free parameters A, b, and c were used for a least-squares fit of ECAP data points between 2.2-2.4 ms and 5.8-6.0 ms. This time range excluded ECAPs from fiber types of interest so that only data points from the decay of the stimulation artifact, and not the neural signal, were fitted. A representative fit is shown in Supplementary Material 7. The fit was subtracted from the raw ECAP trace to produce a detrended ECAP trace, which was used for further quantification of ECAP magnitude.

#### Quantifying ECAP magnitude

A*β*- and B-fiber ECAPs were quantified to investigate their magnitude as recorded by the microneurography, cuff, and LIFE electrodes. A*β*-fibers were selected as a measure of faster-conducting myelinated fibers and B-fibers were selected as a measure of slower-conducting myelinated fibers.

Time windows corresponding to A*β*- and B-fiber ECAPs were calculated for each electrode based on distance from the closest edge of that recording electrode to the center of the stimulation electrode and published values of conduction velocity: 30-70 m/s for A*β*-fibers and 3-15 m/s for B-fibers (Erlanger and Gasser, 1937; Manzano et al., 2008). A tolerance up to 20% was allowed on the conduction velocity if the ECAP was truncated by the initially calculated time window. Then, the fiber-type time windows were further narrowed to be of equal duration across recording electrodes – without removing any portion of the ECAP signal of interest. Equalizing the analysis time window was critical to avoid bias that would arise with increasing distance of the recording electrode from the stimulation electrode. This increased distance translates to a longer computed time window over which the ECAP magnitude calculation is performed – possibly leading to a skewed ECAP magnitude measurement due to the inclusion of additional noise into the measurement. Root mean square (RMS) was calculated on the narrowed fixed-duration window as a measure of ECAP magnitude. RMS was selected over alternative methods (e.g., integral or area under the curve) as it indicates the equivalent steady state energy value of an oscillating signal and represents the noise floor of the recording electrode (Rawlins, 2000). The noise floor is a useful measure, even when no ECAP signal is present at subthreshold stimulation currents. Further, the measure of peak-to-peak voltage is more susceptible to noise and amplitude is more susceptible both to noise and baseline offsets (Mitra et al., 2018).

#### Cohort-based statistics

For ECAP magnitudes and spike counts, the mean value was taken for the replicates of each type of electrode per subject. These mean ECAP magnitudes and spike counts measured on each electrode were normalized to the mean value measured on the cuff contacts and microneurography electrodes, respectively. We performed this normalization for each subject to facilitate comparison across subjects. Log10 was applied to the normalized values to redistribute to normality (Chen, 1994). The Shapiro-Wilk test was used to verify the normality assumption of the student t-tests. One-sided t-tests were used to test the four primary outcomes:

1. The A*β*-fiber ECAP magnitude recorded on the cuff is greater than that recorded on the microneurography electrode
2. The A*β*-fiber ECAP magnitude recorded on the cuff is greater than that recorded on the LIFE electrode
3. The sensory evoked spike count from the microneurography electrode recording is greater than that from the LIFE electrode recording
4. The sensory evoked spike count from the microneurography electrode recording is greater than that from the cuff electrode recording.

Following the pre-registration, *α* = 0.05 was used in a sequential analysis and propagated down the four primary outcomes listed above and then the secondary outcomes until a null result was hit. Confidence intervals (CI) were calculated using the 95% bounds of the t-distribution. Mean is denoted as *µ* and standard error is denoted as (*SE*).

#### ECAP Dose-response curves

Dose-response plots for cVN ECAPs were generated by plotting ECAP magnitude against stimulation current. The A*β*-fiber dose-response curve for each recording electrode was normalized to the ECAP magnitude at 5 mA of stimulation current for that recording electrode, as it was the highest common stimulation current applied across all subjects. Logistic growth functions (sigmoidal shape) were used to fit the dose-response curves following the method of least squares (Castoro et al., 2011). After subtracting the initial offset from the fitted logistic growth functions, EC10 points were calculated for each of the three electrodes for all n=6 subjects. The EC10 point is the stimulation current at 10% of the saturation ECAP magnitude and is a measure of the stimulation level at which the ECAP is first detectable. A repeated measures ANOVA was used to detect if the EC10 point, and hence sensitivity to ECAP detection, was different across the three recording electrodes.

## Results

In this study, we characterized the ability of three commonly used electrodes (i.e., recording cuff, microneurography electrode, and LIFE electrode) to record neural activity from the peripheral nervous system. We investigated both synchronous electrically evoked CAPs and asynchronous spontaneously occurring and sensory evoked neural activity. These results provide the first side-by-side comparison in a human-scaled, large animal model of the recording magnitude, sensitivity, and noise floors of the three recording electrodes in the peripheral nervous system. Further, we compare several referencing strategies and comment on their appropriate use. Lastly, we highlight several artifacts in our recordings that initially appeared to be neural signals but were later classified as artifacts as informed by appropriate control tests. We intend the results of this study to guide the development of neural recording techniques that can be used in the development and deployment of neuromodulation therapies. Direct measures of on-and off-target neural recruitment around the stimulation electrode could advance understanding of therapeutic mechanisms and guide adjustments to electrode design and placement so as to optimize stimulation specificity. Neural activity recordings may also enable continuous closed-loop titration and operation of neuromodulation therapies.

Data associated with this study were shared as part of the Stimulating Peripheral Activity to Relieve Conditions (SPARC) project and are available through the SPARC Data Portal (https://doi.org/10.26275/vm1h-k4kq) under a CC-BY-4.0 license.

### Measuring ECAPs during invasive stimulation

Poor local neural target engagement is critically limiting in some neuromodulation therapies (Heusser et al., 2016; De Ferrari et al., 2017). In these therapies, our ability to directly measure neural activation around the stimulation site and understand fiber types activated could be essential for guiding electrode placement to create more efficacious therapies and better understand those therapies’ mechanisms of action (Verma et al., 2021a). Measurements of local neural target engagement could be done either acutely in the doctor’s office during device placement and therapy titration or chronically by the implanted therapy device. For the acute scenario, a microneurography electrode – already used in clinical research – could be percutaneously introduced to the nerves of interest (Ottaviani et al., 2020). For the chronic scenario, when the treatment is already an invasive device, an additional recording cuff could be implanted on the nerve of interest during the surgery to place the stimulation electrode. Here, we compared the abilities of these two clinically viable electrodes (microneurography electrode and invasive recording cuff) to record ECAPs in the cVN of the domestic swine. Alongside these two devices, we also compared a commonly used pre-clinical electrode, the LIFE electrode (Yoshida and Stein, 1999; Nicolai et al., 2020). Each electrode’s recordings were compared to characterize their ECAP recording magnitudes, sensitivities, and noise floors.

#### Magnitude of ECAP recorded

We characterized the magnitude of the ECAP recorded on each electrode. In general, a larger magnitude signal requires less demanding electronic design to record than a smaller magnitude signal. A representative trace from each electrode is shown in Fig. 3A. ECAPs were recorded concurrently on all three types of recording electrodes (3 contacts each, for a total of 9 recording contacts per subject) at 1.5 mA of stimulation. The stimulation current was selected because it was close to the saturation point of A*β* fibers and provided a stable point of comparison across subjects (n=6).

**Figure 3:**
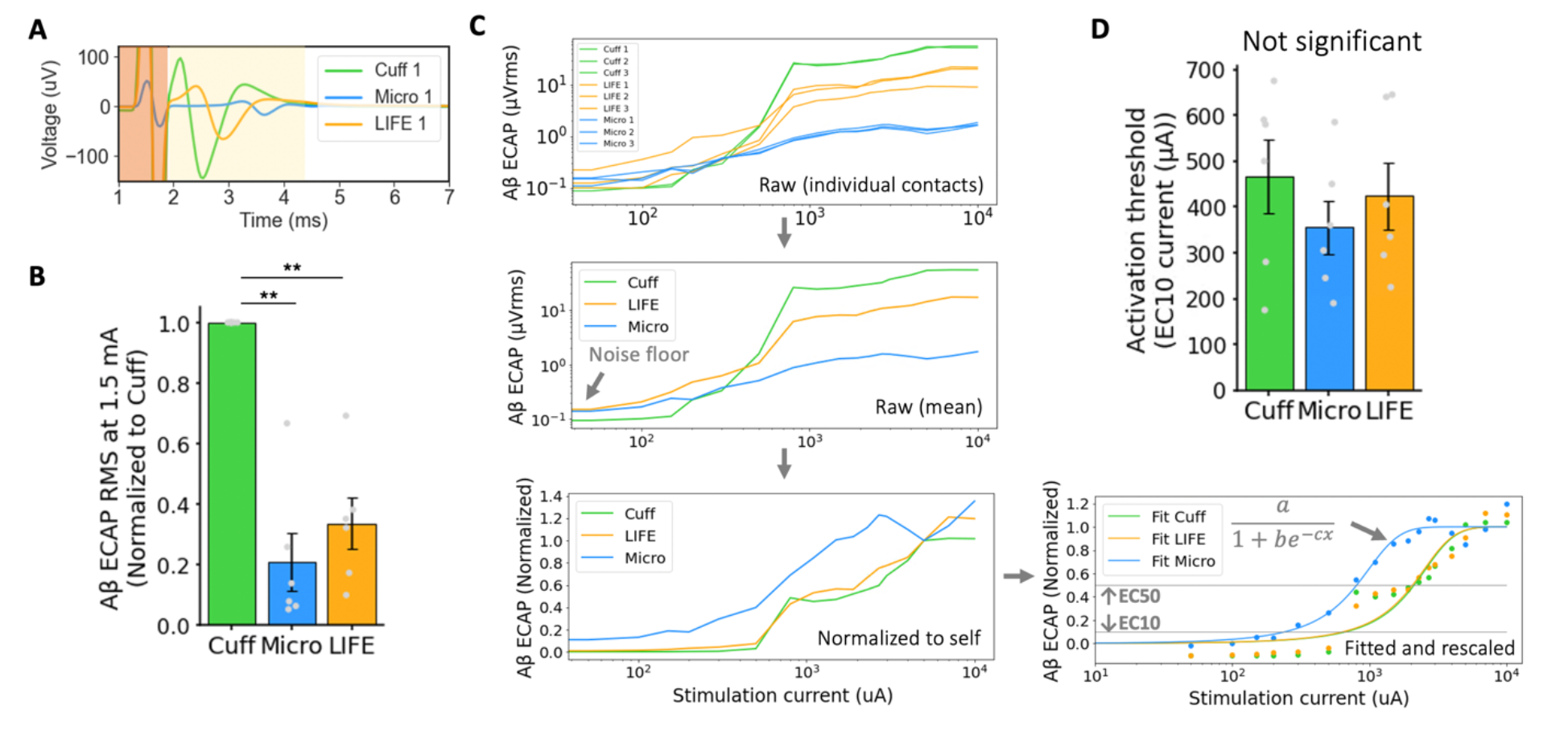
Comparison of electrodes in recording ECAPs from larger myelinated fibers. **(A)** Representative ECAP traces from concurrent cuff, microneurography electrode, and longitudinal intrafascicular electrode (LIFE) recordings at 1.5 mA of stimulation on the cervical vagus nerve (cVN). Contact one of three was selected for each electrode type. Stimulation artifact shaded in orange and A*β*-fiber ECAP shaded in yellow. **(B)** Bar plot summarizing magnitude of A*β*-fiber ECAP recorded with three different electrodes on cVN showing significantly larger recording on cuff compared to both the microneurography electrode and LIFE electrode. **(C)** (top left) Dose-response curves of A*β*-fiber ECAPs for three replicates each of the three recording electrode types characterized. From top left, in an anti-clockwise direction, shows the steps of fitting dose-response curves with logistic growth functions: 1) averaging of replicate contacts, 2) normalization to ECAP amplitude at 5 mA of stimulation, and 3) fitting to logistic growth functions and rescaling y-range to be from zero to one. **(D)** Bar plot summarizing sensitivity of the three recording electrodes across subjects. Sensitivity was measured as EC10 stimulation currents extracted from the dose-response curve logistic growth function fits in **(C)**.

Fig. 3B shows the magnitude of ECAP recorded by each of the recording electrodes during cervical vagus nerve stimulation (VNS). The magnitude of the ECAP recorded by the cuff was significantly larger than the magnitude of the ECAP recorded by the microneurography electrode (normalized to cuff *µ* = 0.208 *SE* = 0.088 *µV_rms_*/*µV_rms_*) (*p* = 0.002, *α* = 0.05, *CI* = [0.052, 0.343]). Similarly, the magnitude of the ECAP recorded by the cuff was significantly larger than the magnitude of the ECAP recorded by the LIFE electrode (normalized to cuff *µ* = 0.335 *SE* = 0.077 *µV_rms_*/*µV_rms_*) (*p* = 0.003, *α* = 0.05, *CI* = [0.147, 0.541]).

#### Sensitivity of recording electrodes

The sensitivity of a recording electrode is critical when the electrode is used to detect the threshold of neural activation, which is a plausible use case in closed-loop control designed to avoid activation of an off-target nerve. The sensitivities of the three recording electrodes were quantified by constructing dose-response curves using 16-18 stimulation current doses between 0-10 mA. The dose-response curves were fitted with logistic growth functions, and the stimulation current at 10% of the maximal ECAP magnitude (EC10) was extracted as a measure of when an ECAP is first detectable (Fig. 3C).

Fig. 3D summarizes the EC10 point of each recording electrode across all subjects (n=6). The microneurography electrode had the lowest mean EC10 stimulation current, suggesting it was the most sensitive recording electrode. However, variability was large and the individual data plots showed that the microneurography electrodes were not consistently the most sensitive electrode in every subject. A repeated measures ANOVA test concluded non-significant differences in EC10 currents across recording electrode type (*p* = 0.26, *α* = 0.05). Repeating the analysis using the EC50 point confirmed a similar trend also with non-significant difference findings (data not shown). Due to this non-significant result, type II error was entirely lost at this secondary outcome and further conclusions in the sequential analysis (see pre-registration) are exploratory, indicated by (*α* = 0) where applicable. Note that the study was powered for the primary outcomes (as reported in the pre-registration) and not this secondary outcome on sensitivity.

Since sensitivity was evaluated by fitting to multiple points in the dose-response curve, it was a measure of the electrodes ‘denoised’ performance and not reflective of the electrodes’ ability to detect the threshold of neural activation from a single level of stimulation current, which depends more critically on the signal-to-noise ratio (SNR).

#### Noise floor of recording electrodes

The electrode’s practical performance in detecting the threshold of neural activation from a single stimulation pulse is determined by the ECAP magnitude and sensitivity of the recording electrode, along with the noise floor of the electrode. The noise floor of each electrode was characterized by calculating the RMS signal across 5 seconds of time series (un-averaged) electrophysiology data both before (Fig. 4A) and after (Fig. 4B) filtering. The noise floor of ECAP traces (Fig. 4C) was characterized by reporting the RMS signal in the A*β* window range at 0 mA of stimulation (no stimulation delivered).

**Figure 4:**
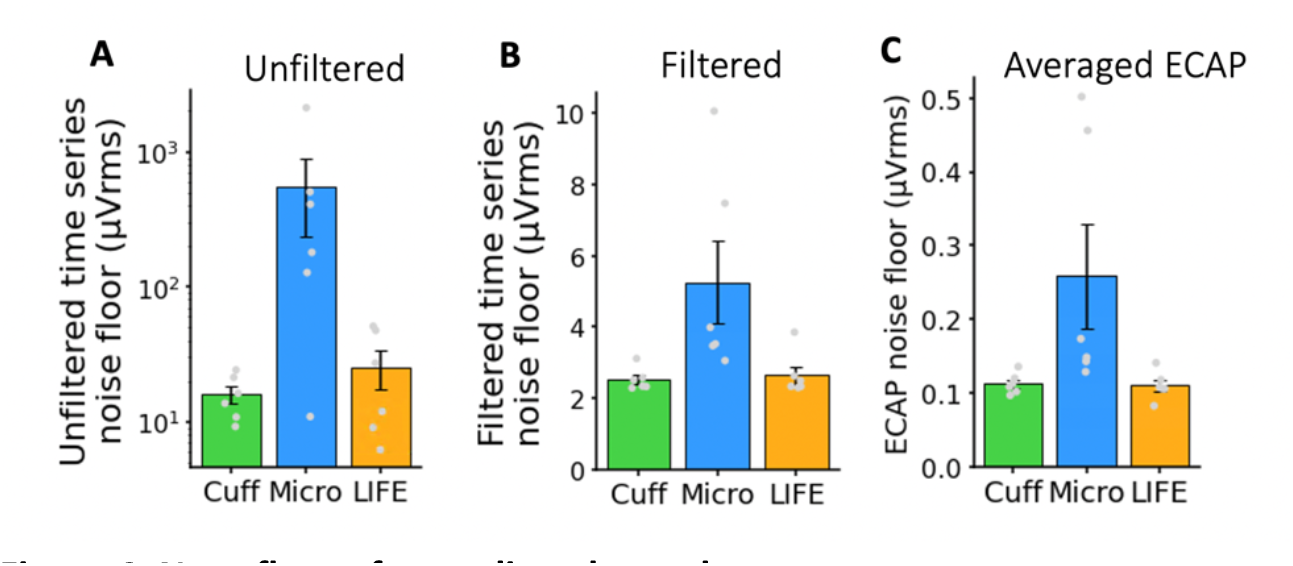
Nose floor of recording electrodes. Noise measurements in electrophysiology recordings showing largest noise levels on the microneurography electrode. Note the different y-axis scales in the three plots as the noise level after filtering and averaging is reduced by orders of magnitude. **(A)** Unfiltered time series electrophysiology. **(B)** Filtered time series electrophysiology. **(C)** Filtered averaged ECAPs (median of n=750).

,H3>B-fiber recordings on recording electrodes

Fig. 4 shows the noise floor of all three recording electrodes before and after filtering in time series and after filtering in averaged (n=750) ECAP traces. As expected, the noise floor is higher in the time series and lower in the averaged ECAP traces (2.53 *µV_rms_* vs. 0.11 *µV_rms_* taking the recording cuff as an example). In the time series electrophysiology data, the noise is higher before filtering than after (15.85 *µV_rms_* vs. 2.53 *µV_rms_* taking the recording cuff as an example), largely due to the contribution of low frequency noise (Supplementary Material 8). The microneurography electrode has the highest noise floor across all three electrodes (0.26 *µV_rms_* for the microneurography electrode vs. 0.11 *µV_rms_* for the cuff and LIFE, taking the filtered ECAP traces as an example). In subject 5 and 6, an active head stage was used on the microneurography electrode, which pre-amplified the signal at the electrode, theoretically reducing SNR loss due to noise pick up on the wire from the electrode to the main amplifier. However, in using the active head stage, additional electronic noise was introduced by the pre-amplifier and is observed in the two higher noise data points in Fig. 4B (mean of 3.51 *µV_rms_* for subjects 1-4 vs. 8.77 *µVrms* for subjects 5-6, taking the filtered time series as an example). We summarize and compare the sources of noise (i.e., environmental, biological, electrochemical, and electronics) between the recording electrodes in the Discussion section.

Smaller diameter myelinated fibers are hypothesized to be responsible for the therapeutic effects of several neuromodulation therapies. For example, B-fibers are putatively responsible for the therapeutic effects in the heart failure application of VNS (Sabbah et al., 2011). B-fibers are more challenging to record from due to the smaller fiber size and signal magnitude (Yoo et al., 2013; Nicolai et al., 2020). We compared the three recording electrodes in their ability to record B-fiber ECAPs from the cVN. Vecuronium was administered at a sufficient dose in subjects 4-6 to blunt muscle activity and prevent EMG artifact from contaminating the neural recordings. However, due to a protocol restriction, insufficient vecuronium was administered in subjects 1-3 to fully prevent EMG artifact from contaminating the neural recordings. B-fibers were differentiated from EMG artifact by verifying signal propagation delay across the replicate contacts of each electrode type (Fig. 5A, inset). It was not possible to conclude that we were quantifying only B-fibers because of the overlap of B-fibers and A*δ*-fibers in the conduction velocity ranges (Erlanger and Gasser, 1937; Manzano et al., 2008). However, the purpose of this analysis was only to contrast slower-conducting myelinated fibers (B/A*δ*) with the previous, faster-conducting myelinated fibers (A*β*). Analyzing results from two distinct fiber types allowed us to investigate any differences in ECAP magnitude across the recording electrodes by fiber type and hence widen applicability of the results.

**Figure 5:**
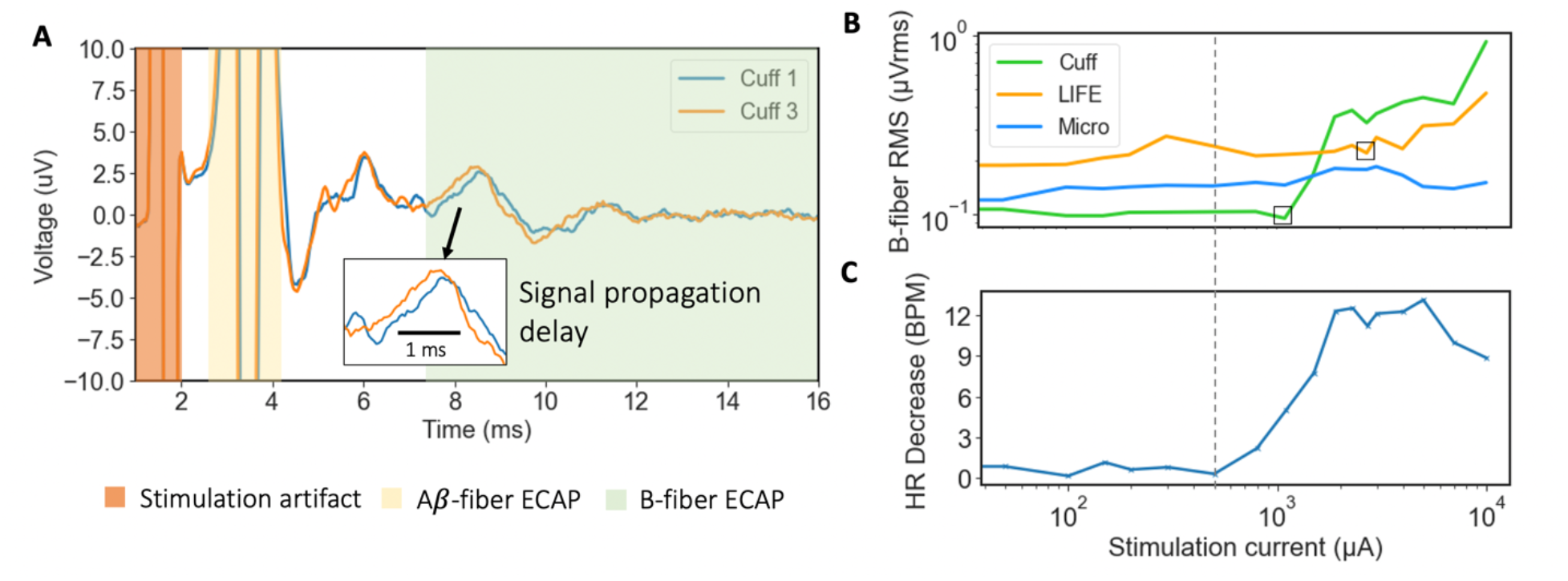
Comparison of electrodes in recording ECAPs from smaller myelinated fibers and correlation to evoked bradycardia. **(A)** Authenticity of B-fiber recordings was confirmed by signal propagation delay across recording electrodes in the range of 3-15 m/s (Erlanger and Gasser, 1937; Manzano et al., 2008). An artifact (e.g., EMG, motion) could occur in a similar time window but would not show up with a signal propagation delay across spatially separated recording electrodes. Stimulation artifact shaded in orange, A*β*-fiber ECAP shaded in yellow, and B-fiber ECAP shaded in green. Stimulation was delivered at 10 mA. A similar plot is shown for the LIFE and microneurography electrodes in Supplementary Material 9. **(B)** B-fiber dose-response curves for all three recording electrodes on a single subject showing the largest ECAP magnitude recorded with the cuff electrode. The cuff electrode also recorded ECAPs at the lowest stimulation current. The ECAP detection threshold for each electrode is boxed; a clear threshold was not apparent on the microneurography electrode. The dash line from **(C)** indicates the bradycardia onset threshold. **(C)** Evoked heart rate decrease (bradycardia) dose-response curve in the same subject as **(B)**. The bradycardia stimulation current threshold is lower than the B-fiber ECAP detection thresholds in **(B)**.

B-fiber ECAPs were most consistently recorded on the cuff, only evident in certain subjects with the LIFE electrode, and not evident with the microneurography electrode. Fig. 5A shows B-fiber ECAPs confirmed by signal propagation delay across two cuff contacts. Similar plots to Fig. 5A for the other electrodes show authentic B-fiber ECAPs recorded on the LIFE electrode but not the microneurography electrode (Supplementary Material 9). Fig. 5B and 5C show doseresponse curves for B-fiber ECAPs and stimulation-evoked bradycardia, respectively. B-fibers are putatively responsible for decreases in heart rate during VNS (Qing et al., 2018). The bradycardia dose-response curve best tracks the cuff B-fiber recordings up to ~2 mA. After that, afferent pathways may be recruited counteracting the efferent B-fiber mediated effects (Ardell et al., 2017). The cuff B-fiber ECAP recordings correlate best with stimulation-evoked bradycardia in this subject (R_cuff = 0.70 and R_LIFE = 0.39). Data from remaining subjects are shown in Supplementary Material 10.

### Reference electrode on nerve versus in surrounding non-neural tissue

Judicious choice of reference electrode placement can optimize neural signal fidelity and minimize artifacts (Sabetian et al., 2017). Neural recordings can be referenced to a location on the active nerve itself or in local non-neural tissue. Conventionally, cuff electrodes have been referenced on the nerve (e.g., bipolar or tripolar), while in microneurography, the reference is placed in local non-neural tissue (Macefield, 2021). We conducted ECAP recordings with different referencing strategies and systematically compared ‘on-nerve’ references with ‘local tissue’ references.

Fig. 6A shows that local tissue referenced recordings have a much larger stimulation artifact than on-nerve referenced recordings, but also a ~2.5x larger peak-to-peak ECAP magnitude. The smaller magnitude ECAP recorded with on-nerve references may be explained by ‘subtraction’ of neural signal between the recording and reference electrode due to the 2-3 cm wavelength of the A*β*-fibers that appears common to both electrodes (Andreasen and Struijk, 2002). The ECAP can be visualized as a moving electric dipole with the negative depolarizing charge leading the positive repolarizing charge (Fig. 1B). The distance between the poles is the ‘wavelength’ of the ECAP and positively relates to fiber diameter. For smaller fibers, with wavelengths less than the distance between the recording and reference contacts on the nerve, this subtraction effect should not be evident. Thus, on-nerve references may be superior to local tissue references when recording smaller diameter fibers, as ECAP magnitude may be preserved while common mode artifact is reduced.

**Figure 6:**
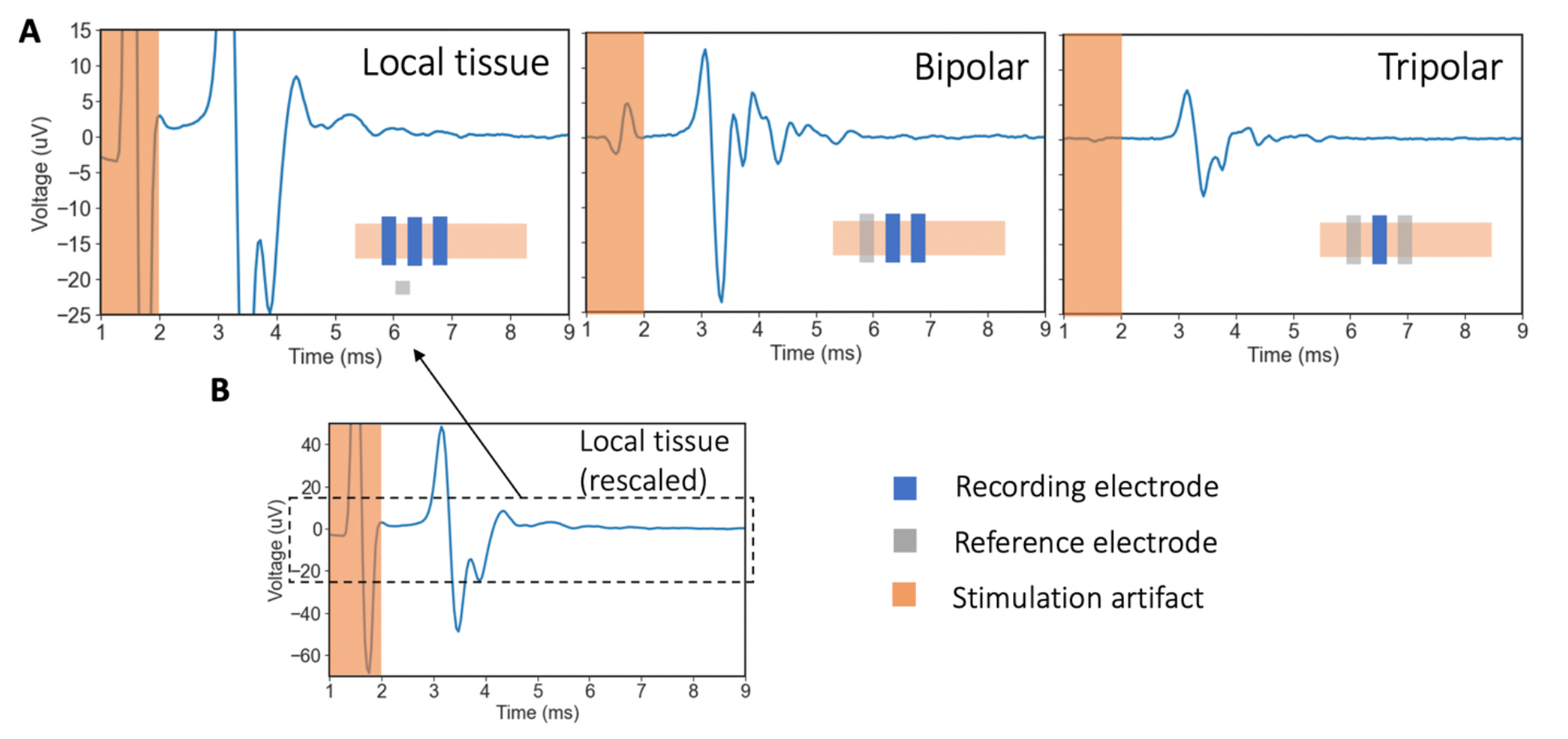
Effect of referencing strategy on stimulation artifact and ECAP. **(A)** Representative data showing ECAP traces with three different referencing strategies at 2.3 mA of stimulation: (left) reference in ‘local tissue’, (center) bipolar ‘on-nerve’ reference, and (right) tripolar on-nerve reference. Orange highlights overlap the stimulation artifact, an example of common mode noise, which is substantially reduced in the on-nerve referencing strategies. The A*β*-fiber ECAP (~3-4 ms) is also smaller in the on-nerve referencing strategies. In the illustration of the recording setup in each plot, recording electrodes are in blue and reference electrodes are in gray. **(B)** Zoomed out plot of local tissue referenced recording from **(A)** to show larger y-axis range.

#### Tripolar versus bipolar on-nerve referencing strategies with recording cuff

An on-nerve reference close to the active recording electrode can efficiently reduce common-mode artifacts external to the electrodes, such as spillover electric fields from muscle activity and stimulation artifact (Sabetian et al., 2017). Cuff recording electrodes lend themselves to on-nerve reference strategies and are typically employed in a bipolar or tripolar configuration. Past studies have shown several key differences between tripolar (contact 1 and 3 tied together to reference, contact 2 used for recording) and bipolar referenced cuff recordings (Sabetian et al., 2017). Namely, tripolar referencing reduced the representation of common mode artifacts (Sabetian et al., 2017), which is pertinent because these artifacts can appear at similar time points as neural fiber signals (Nicolai et al., 2020). However, bipolar cuffs, compared to tripolar cuffs, can detect neural signal propagation direction (afferent vs. efferent). Meanwhile, the magnitude of neural signal recorded by the bipolar vs. tripolar cuff depends on the geometry of the cuff. In particular, the ECAP magnitude recorded depends on the electrode length (EL) and electrode edge length (EEL) of the insulation, which are further explored in computational work by Sabetian and colleagues (2017).

We collected both tripolar and bipolar referenced recordings using the same cuff. Fig. 6A shows representative bipolar and tripolar traces at 2.3 mA of stimulation. The stimulation artifact magnitude is smaller with the tripolar compared to the bipolar reference. The peak-to-peak ECAP magnitude is also smaller with the tripolar reference by a factor of ~2.2x compared to the bipolar reference. Data from other subjects are shown in Supplementary Material 10 and support the conclusions reported here. Our experimental results support the computational results reported by Sabetian and colleagues (2017), which predicted a ~1.4x larger peak-to-peak neural signal on the bipolar compared to the tripolar referenced recordings (EL = 9 mm and EEL = 2 mm).

### Measuring asynchronous neural activity

Changes in spontaneously occurring neural activity could be a sensitive and real-time marker for neuromodulation therapies (Gonzalez-Gonzalez et al., 2021; Chao et al., 2021). Measuring spontaneous activity may be especially important in cases when there is an afferent-efferent pathway with one or more synaptic connections, which introduces variable conduction latency and desynchronizes the timing of the neural signal, thereby preventing a clear ECAP from appearing in the later efferent pathway. Instead, changes in spontaneously occurring neural activity of the later efferent pathways can be a sensitive measure of therapy effect and can be used for therapy titration. Recording spontaneous activity may also be important for non-traditional waveforms such as burst, pink noise, high frequency block, or direct current (DC) block wherein the timing of initiation of an action potential may be variable or moot, and the intended effect is to change spontaneous activity in between stimulus trains. Consequently, we compared the three recording electrodes in their ability to measure sensory evoked neural activity, an asynchronous neural signal, from the great auricular nerve (GAN), which is implicated as an on-or off-target nerve in various auricular stimulation-based therapies.

Fig. 7B shows representative data of sensory evoked neural activity recordings from a cuff electrode, microneurography electrode, and LIFE electrode during a 30 second quiescent period, 30 seconds of on-target stroking at a region of the auricle innervated by the GAN, and 30 seconds of off-target stroking at a region of the auricle not innervated by the GAN (Fig. 7A). The representative recording from the cuff electrode showed spike-like activity throughout the 90 second recording period. The shape of the spike (Fig. 7C, bottom panel) and the synchronization of the timing with the ventilator indicated that the spike activity on the cuff electrode was due to motion artifacts from the ventilator. Transection of the GAN was also attempted as a control to further verify the authenticity of the neural recordings. However, due to the webbed nature of the nerve at the insertion point to the ear, we were unable to completely transect it in every subject to eliminate the sensory evoked neural activity.

**Figure 7:**
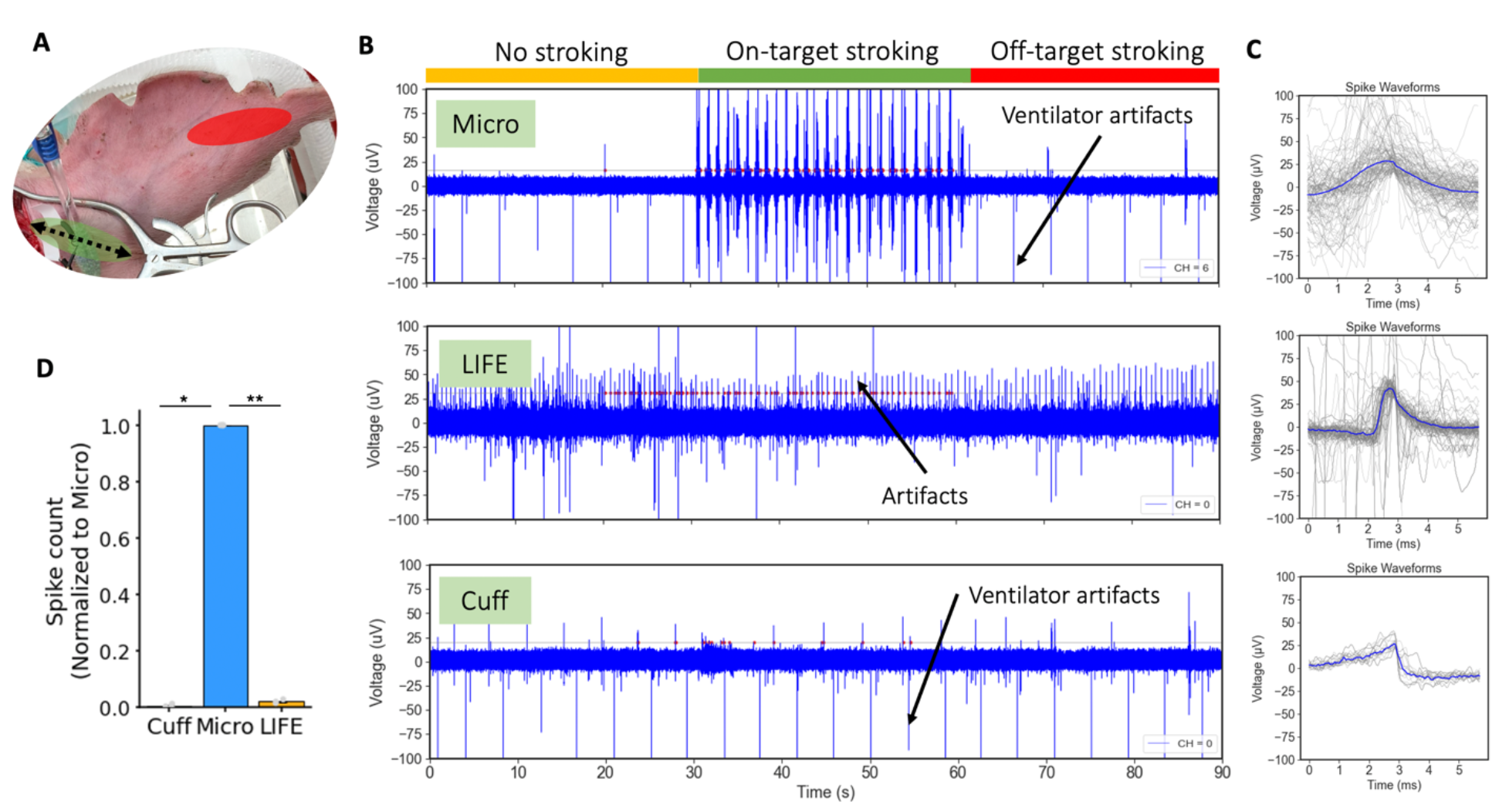
Comparison of electrodes in recording asynchronous sensory evoked neural activity. **(A)** Using a toothbrush to gently stroke the ear at off-target (red, not innervated by GAN) and on-target (green, innervated by GAN) areas corresponding to green and red color coding in **(B)**. **(B)** Representative electrophysiology recordings: first 30 seconds with no stroking, middle 30 seconds with on-target stroking, and last 30 seconds with off-target stroking to control for motion artifacts from stroking. Only the microneurography electrode showed a robust response specific to on-target stroking. LIFE and cuff recordings were selected to illustrate artifacts, which appeared neural, but were not authenticated by appropriate controls. The gray line in each plot is the spike detection voltage threshold and the red dots indicate detected ‘spikes’. **(C)** Spike shapes for electrophysiology recordings in **(B)** indicating multi-unit hash recordings on the microneurography electrode, tremor or cardiac motion artifact on the LIFE electrode, and ventilator artifact on the cuff electrode. **(D)** Bar plot summarizing spike count recorded by the three electrodes across all subjects showing the microneurography electrode records significantly more spikes than both the cuff and LIFE electrodes.

The bar plot in Fig. 7D summarizes the spike count on each recording electrode (normalized to the microneurography electrode in that subject) during on-target sensory stroking of the region of the auricle innervated by the GAN. The microneurography electrode recording spike count was significantly larger than the cuff recording spike count (normalized to microneurography *µ* = 4.38 × 10−3 *SE* = 2.43 × 10−3) (*p* = 0.01, *α* = 0.05, *CI* = [5.22 × 10−5, 6.34 × 10−2]). Similarly, the microneurography electrode recording spike count was significantly larger than the LIFE electrode recording spike count (normalized to microneurography *µ* = 1.95 × 10−2 *SE* = 2.7 × 10−3) (*p* = 0.001, *α* = 0.05, *CI* = [9.38 × 10−3, 3.76 × 10−2]).

Spontaneous activity recordings collected from the cVN did not show authentic neural spikes with any of the recording electrodes (Supplementary Material 11). Signals that initially appeared neural were likely motion artifacts as they persisted more than 20 minutes after the nerve was double transected cranial and caudal to the recording electrodes. Further, the artifacts were more pronounced in subjects 1-3, where the subjects experienced more tremor compared to subjects 4-6.

### Artifacts in electrophysiology recordings

We applied several controls to differentiate artifacts from bona-fide neural signals. These included: administration of a muscle paralytic, nerve transections, verifying signal propagation delay across recording contacts, concurrent monitoring of heart rate and respiration to investigate synced ‘neural’ activity, inspection of waveform shapes, and controls for induced motion. With these controls, we classified several signals that initially appeared neural as non-neural artifacts. Specifically, in time series (non-averaged) recordings we identified artifacts due to respiration, cardiac, and tremor effects.

Firstly, respiration related artifacts are captured in Fig. 7B (cuff recording) occurring at a rate of once every ~4s (15 times/minute) in sync with the respiration rate set on the ventilator. The artifact was still present after nerve transections and had a unique non-natural spike shape (Fig. 7C, bottom panel versus sensory evoked activity in top panel). The ventilator-induced artifacts likely occurred due to motion of the animal with respiration. Secondly, tremor-related effects are observed in Fig. 7B (LIFE recording). Subject tremor is described in the Methods. These motion related artifacts from tremors disappeared with the administration of a muscle paralytic, vecuronium, or the anesthetics Telazol or Ketamine, all of which stopped the tremors. Lastly, cardiac related artifacts were recorded (shown in Supplementary Material 8), which were removed with high pass filtering at 100 Hz. Cardiac artifacts can be directly from electrocardiogram (EKG) electrical activity or from cardioballistic effects due to the beating motion of arteries adjacent to the nerve (Ulrich et al., 2016), which causes motion in the nerve and the recording electrode. The cardioballistic effect was visible to us in several instances and was synced with heart rate. Direct EKG may be a more likely source of the artifact if the recording and reference electrode pair are across the heart or generally separated by a larger distance. The susceptibility of all three recording electrodes to motion induced artifacts suggests that one needs to be careful in interpreting spontaneous neural activity recordings in response to physiological events that also create motion, such as respiration, cardiac activity, etc.

## Discussion

In the Discussion, we first summarize the strengths and weaknesses of different recording electrodes and discuss suitable electrode type and referencing strategy for each application. Second, we outline potential mechanisms of neural recording to explain the results. Third, we discuss challenges in recording spontaneously occurring activity and TENS evoked CAPs. Fourth, we discuss sources of noise in electrophysiology recordings. Finally, we discuss correlation of ECAP magnitudes recorded on the three electrodes with evoked cardiac responses.

### Electrode choice by application

In the context of neuromodulation therapies, ECAPs can be used as a real-time and precise measure of neural target engagement around the stimulation electrode (Verma et al., 2021a). Several neuromodulation therapies are critically limited by off-target nerve activation (Heusser et al., 2016; De Ferrari et al., 2017), which may be addressed by electrode design and stimulation parameters. Electrode design and stimulation waveforms are often translated directly from animal models and not adjusted for human use due to lack of real-time measures to guide development. We believe ECAPs represent a readily available method to guide development and deployment of neuromodulation therapies to improve on-target neural engagement and consequently therapeutic efficacy. Table 1 summarizes the performance and utility of the different recording electrodes based on the outcomes of the experiments performed in this study.

**Table 1:**
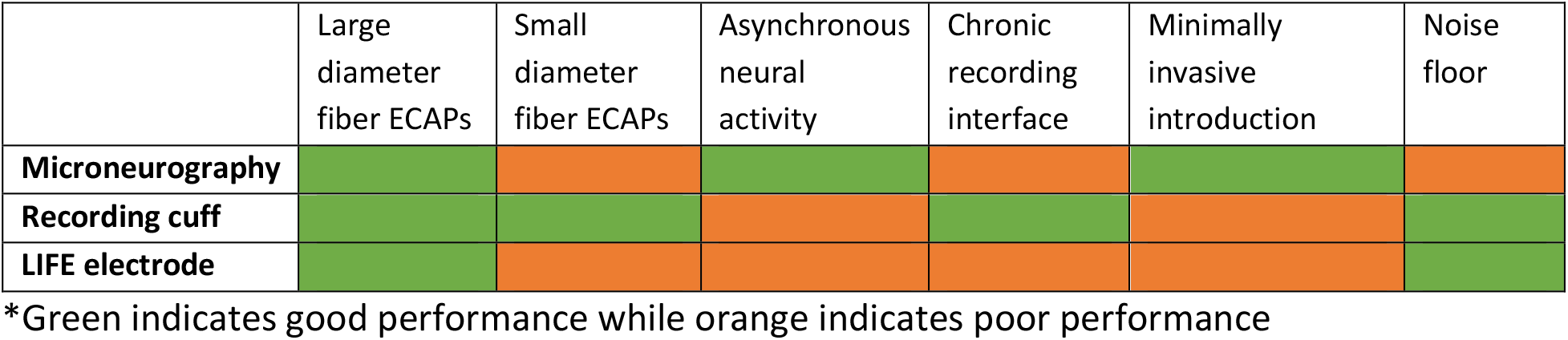
Recording electrode choice by application

#### Use of microneurography electrodes to develop and deploy minimally invasive neuromodulation therapies

Our data showed that recordings made using microneurography electrodes consistently measured A*β*-fiber ECAPs. In fact, once full ECAP dose-response curves were constructed and fitted to overcome the SNR limitations of the microneurography electrodes, they showed the highest sensitivity in detecting an ECAP amongst the three recording electrodes tested. We also showed that recordings made using microneurography electrodes most consistently measured sensory evoked neural activity compared to recordings made using cuff and LIFE electrodes.

The microneurography electrode tested is already used in humans to record muscle sympathetic nerve activity (MSNA) from the common peroneal nerve (Macefield, 2021) and median nerve (Macefield, 2021) and, most recently, to record neural activity from the cVN (Ottaviani et al., 2020). Similar to this existing human use, a microneurography electrode may be inserted percutaneously during clinical trials and at the deployment of neuromodulation therapies to map sensory innervation (Meier et al., 2018) and inform non-invasive stimulation electrode placement. The temporarily inserted microelectrode can also measure stimulation evoked on- and off-target neural activation, and that data can be used in real-time to adjust stimulation electrode position, design, and stimulation parameters (Verma et al., 2021a).

To aid in the translation of the microneurography technique to routine clinical use, the technique needs to be easy to perform and deployable within minutes. Currently, the technique requires a skilled microneurography practitioner, and even then, it can be time consuming for that practitioner to target specific fascicles in a nerve. To overcome these obstacles, automation, such as insertion based on force or electrical impedance (Kodandaramaiah et al., 2012; Park et al., 2018), can be used to guide the microneurography electrode into the nerve and specific fascicle, where necessary. The epineurium and perineurium have vastly different mechanical properties and may be ideal for differentiating between fascicles (Koppaka et al., 2022).

#### Use of cuff electrodes to provide high-fidelity chronic recordings

Our data showed that the cuff electrode recorded ECAPs with the highest SNR. The cuff was also the only electrode to consistently record smaller diameter myelinated fiber (e.g., B-fiber) ECAPs. An implanted recording cuff would be a suitable option in applications where a chronic recording interface is necessary and smaller fiber types are of interest.

The high SNR of the cuff ECAP recordings will be advantageous in closed-loop strategies that rely on detecting first activation of a particular fiber type, and where time for averaging is limited. A potential application could be a waveform block strategy attempting to block larger diameter fibers while activating smaller diameter fibers. In this application, sensitive and realtime monitoring of the large diameter fiber activity will be required to ensure that the nerve block is sustained. If any large diameter fiber activity is detected, the stimulation parameters must be quickly adjusted to restore the nerve block. The recording cuff provides the highest SNR ECAP recording from a single stimulation pulse, without the time delay of averaging, required by such applications.

### Referencing strategy

The results show that referencing strategies (i.e., on-nerve bipolar vs. on-nerve tripolar vs. local non-neural tissue) have a profound effect on recorded ECAP and artifact magnitude (Fig. 6). On-nerve references reject common mode noise more effectively but also abate the recorded ECAP magnitude. The microneurography technique lends itself to the local tissue reference (Macefield, 2021), resulting in larger magnitude ECAPs recorded, which compensates for the lower ECAP magnitudes otherwise observed with the microneurography electrode. To minimize common mode noise, the microneurography electrode should be inserted approximately equidistant from the stimulation site as the recording electrode and in fat. The reference electrode tip should also be of similar material and design to the recording electrode to match the representation of common mode artifact (Ludwig et al., 2008). The cuff electrode lends itself to on-nerve referencing strategies, which reduces the magnitude of the ECAPs recorded (Fig. 6A), due to ‘subtraction’ of the neural signal when it appears common on both the recording and on-nerve reference electrode. See Results section for a discussion on how the ‘subtraction’ effect is less prominent for smaller fiber types (with shorter wavelengths) and with larger spacing between the cuff recording and reference contacts. We observed that the cuff otherwise recorded the largest magnitude ECAPs, thereby balancing out this ‘subtraction’ effect due to on-nerve referencing. The on-nerve reference strategies performed better at rejecting common mode artifacts, which is critical in out-of-clinic ambulatory settings where a chronically implanted cuff would likely be required regardless. Therefore, we conclude that local tissue reference, which is conveniently implemented in microneurography recordings, and on-nerve reference, which is conveniently implemented in cuff recordings, are appropriate for use as such.

### Mechanisms of neural recording

Surprisingly, although recordings made with the microneurography electrode showed the most distinct sensory evoked neural activity (Fig. 7B), they measured A*β*-fiber ECAPs with the smallest magnitude (Fig. 3 A-B). In contrast, the recordings made with the cuff did not show distinctive sensory evoked neural activity but measured ECAPs with the largest magnitude. The sensitivity of the penetrating microneurography electrode may be explained by its proximity to the firing neural source. The larger ECAP magnitude on the recording cuff may be explained by the cuff more linearly summing the electric potentials from all neurons firing within it (Humphrey and Schmidt, 1990). On the other hand, the microneurography electrode sums up potentials with a greater weight from neurons firing adjacent to the recording tip and a substantially reduced weight from neurons at a distance from the recording tip (Buzsáki, 2004; Khodagholy et al., 2015). In this manner, the microneurography electrode records a large signal from an adjacent neural source and a small signal from a further neural source, while the cuff records and sums moderate signals from throughout the nerve it is applied on.

### Recording spontaneous neural activity

None of the recordings from the cVN showed clear and repeatable spontaneously occurring neural activity (Supplementary Material 11). This contrasted with the clear recordings of sensory evoked activity from the GAN recorded by the microneurography electrode. This also contrasted with past microneurography work, which showed spontaneous activity recordings from the cVN of humans (Ottaviani et al., 2020). Compared to the human recordings, which were done percutaneously, we had an open surgical preparation. This may have led to additional trauma during electrode insertion with less mechanical stability provided by surrounding tissue. Further, the largest spontaneous signals from the cVN are expected to come from A*β* muscle afferents under muscle tone. The manipulation of these muscles and the application of the muscle paralytic, vecuronium, in our experiments may have substantially reduced neural signaling from these muscles compared to clinical recordings in awake subjects. Similarly, the sensory evoked neural activity on the GAN is expected from A*β* fibers, which are a large fiber type and produce larger extracellular signals than the smaller B and C fibers, which may be the only source of spontaneous activity remaining on the cVN under the mentioned surgical and anesthetic conditions.

Given our results, it is surprising that experiments in rodents have shown recordings of spontaneous activity from the cVN with simple hook electrodes (similar to cuff but without insulation) (Silverman et al., 2018; Stumpp et al., 2020). The difference in nerve size and perineurium thickness between rodents and pigs or humans may explain these results (Koh et al., 2022). As illustrated in Fig. 1C, the neurons in a rodent are much closer to the recording cuff than in a pig or human. The shorter electrode-fiber distances in rodents results in a stronger extracellular signal at the point of the recording cuff and larger amplitude neural activity recordings (Koh et al., 2022).

While clear sensory evoked neural activity was not evident on the macroelectrode (cuff and LIFE electrode) recordings on the GAN, it is possible that the neural signals were masked by the low SNR of the recordings. Metcalfe and colleagues (2018) have shown that velocity selective addition of neural signals from multiple spatially separated channels, can be used to improve the SNR and detect spontaneous neural activity in macroelectrode recordings of human-scaled peripheral nerves. Such strategies may be considered if the conduction velocity of the fiber of interest is known and computational resources to implement the algorithms are not limited by the intended clinical application. The absence of spontaneous activity in recordings collected using macroelectrodes compared to microelectrodes is also seen in the central nervous system and in humans (Winestone et al., 2012).

### Measuring ECAPs during non-invasive stimulation

Numerous non-invasive stimulation devices have hit the market in recent years. Several claim to stimulate nerves deep below the surface of the skin without activating superficial off-target nerves (Nonis et al., 2017; Verma et al., 2021b). Microneurography-based studies (Ottaviani et al., 2020) that attempt to record non-invasive ECAPs from the intended on-target nerve and expected off-target nerves during delivery of the non-invasive stimulation could clarify the actual nerves responsible for effect and side effect in an already efficacious therapy. Then, in future deployment of the therapy, the same microneurography techniques can be used to customize electrode position to account for inter-subject variability in anatomy. Customizing electrode position to the individual’s anatomy will ensure that the same on-target nerve is recruited in each patient – possibly increasing responder rates. Direct and real-time measures of local neural target engagement are critical, as surrogate or indirect measures can be confounded by muscle activity (Feucht and Ward, 2021), poor temporal resolution, and other factors (Verma et al., 2021a).

We used TENS electrodes (2 × 2 cm^2^) to stimulate 1) the region of the ear innervated by the GAN and 2) the base of the ear through which the trunk of the GAN courses. Using up to 10 mA of stimulation current, we did not detect ECAPs with any of the three recording electrodes (data not shown). While this current density was comparable to the lower end of current densities used in aVNS clinical trials (Verma et al., 2021a), and should have been sufficient for perception, it may have been insufficient to create enough synchronous recruitment of neurons to record clear ECAPs. The larger surface area of the TENS electrodes translates to temporally synchronous activation of fibers but at spatially distinct locations across the span of the TENS electrode. As the neural signals propagate, they remain asynchronous and do not form a clear large magnitude ECAP at the recording electrode. Janko and Trontelj (1980) reported on microneurography recordings from the median, radial, and ulnar nerve during non-invasive stimulation of the finger. They used a smaller surface area stimulation electrode, which would have lent to more synchronized recruitment of neurons and a clearer ECAP. These findings may have implications towards stimulation electrode design for non-invasive neuromodulation therapies – especially if the mechanism of action is reliant on synchronous or coordinated recruitment of neurons.

### Sources of noise in electrophysiology recordings

Fig. 4 summarizes the noise floor of each recording electrode before and after filtering and with averaging for ECAP presentation. Several key sources of noise in neural recordings are listed here in order from the external environment, the *in vivo* environment, the electrode, and finally the electronics:

- External noise capacitively and/or inductively coupled, conducted, or radiated (in the case of higher frequency electromagnetic field), into the recording, including mains 50/60 Hz noise (Yasar, 2021).
- Biological noise consisting of background neural and other electrical activity in the body (Yang et al., 2009).
- Noise due to motion of the electrode relative to tissue (Simakov and Webster, 2010).
- Thermal noise, generated by thermal motion of charge carriers and dependent on electrode impedance (Yang et al., 2009).
- Electrochemical noise reflecting corrosion on the metal electrode and interaction of charged ions with the solution (Obot et al., 2019).
- Electronic noise in the recording equipment including flicker noise, thermal noise, digitization noise, etc. (Molecular Devices, 2006).

We found that electronic noise dominated in the 0.1 – 3 kHz filtered time series relevant to peripheral ECAP and spiking activity. We measured the electronic noise of the TDT SIM head stage to be 1.95 *µVrms*. The measurement was made by shorting the input to reference while the setup was in a Faraday’s cage (to reduce pick up of external noise). The recorded time series trace was then filtered, and RMS noise was calculated as it was for *in vivo* recordings (Fig. 4). Our measurement of the input referred noise floor was verified by the TDT SIM head stage datasheet values after adjustment for the appropriate bandwidth (TDT, 2022). *In vivo* measurements of noise from the recording cuff filtered time series trace was 2.53 *µVrms* on average (Fig. 4B), which suggests that ~77% of RMS noise is from the recording equipment electronics front end and digitization. This noise is highly uncorrelated: averaging over 750 pulses, to construct ECAP trace presentations, reduces noise on the recording cuff to 0.11 *µVrms* compared to a theoretical reduction by 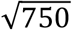 to 0.09 *µVrms* if the noise was entirely uncorrelated (Metcalfe et al., 2018). Since most of the noise in this recording frequency band is coming from electronics front-end and digitization, application specific front-end and analog to digital converter (ADC) architecture and design can be exploited to reduce noise. Further tightening the recording bandwidth will also reduce RMS noise. Note that electronic noise may not be the dominant noise source in other recording frequency bands, such as local field potential (LFP) recordings.

For the higher impedance microneurography electrode, we see additional noise contributions for a total noise floor of 3.51 *µVrms* in the microneurography electrode filtered time series trace versus 2.53 *µVrms* in the lower impedance recording cuff trace. This difference of 0.98 *µVrms* is likely attributable to increased electrode thermal noise, increased biological noise from background neural activity, and/or increased external noise pickup due to the higher impedance of the microneurography electrode. Note that although the impedance of the microneurography electrode was specified as 1 MOhm at 1 kHz, 1 MOhm would not be used as the resistance (*R*) to calculate thermal noise in the equation (Molecular Devices, 2006):

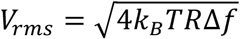

This is because the 1 MOhm includes both real and imaginary impedances. Only real impedance (resistance, *R*) is used to calculate thermal noise. An ideal capacitor, despite having an imaginary impedance, does not generate thermal noise (Lundberg, 2002). The true thermal noise of the electrode is the noise calculated with the real impedance and filtered by the equivalent circuit capacitance of the electrode (Lundberg, 2002). The true thermal noise of the microneurography electrode would be much lower than the 6.9 *µV_rms_* calculated incorrectly using *R* = 1 MOhm. This is supported by the experimental results where there was only an additional 0.98 *µV_rms_* noise on the much higher impedance (~1 MOhm) microneurography electrode compared to the lower impedance (~1 kOhm) cuff electrode.

Our results also showed that all three recording electrodes were susceptible to motion induced artifacts from respiration, cardiac activity, tremoring, etc. In this animal study, we were able to use controls such as nerve transections and administration of a muscle paralytic, vecuronium, to differentiate neural signals from motion artifacts. However, use of these controls may not be feasible in human studies. Therefore, practitioners need to be careful in interpreting clinical microneurography recordings that are expected to correlate to physiological events that also create motion, such as respiration, cardiac activity, etc. Inspection of neural spike shapes and control recordings from close-by, non-neural tissue can help differentiate between neural signals and artifacts. High pass filtering to remove lower frequency artifacts and placement of the reference electrode close to the recording electrode to lower common mode artifacts can further reduce motion induced artifacts in the recordings.

### Correlation of VNS cardiac response and B-fiber ECAPs

A previous publication from our group showed strong tracking between A*a*-fiber ECAP recordings and EMG activity recorded from the laryngeal muscles, implicated as off-target activation in clinical VNS for epilepsy (Blanz et al., 2022). However, the publication did not investigate tracking between stimulation-evoked bradycardia and B-fiber ECAPs, which are putatively responsible for bradycardia during VNS (Qing et al., 2018).

In Fig. 5, we show correlation between stimulation-evoked bradycardia and B-fiber ECAPs magnitude from a select subject. The correlation was strong with cuff ECAP recordings (R_cuff = 0.70) but poor with LIFE recordings (R_LIFE = 0.39), which were the electrodes used exclusively in the previous study by our group (Blanz et al., 2022). Interestingly, shown in Fig. 5C, we saw an increasing bradycardia response up to a VNS amplitude of ~2 mA, and then a flat and subsequently a decreased bradycardia response. We hypothesize that at higher stimulation currents, additional fiber types, which have a tachycardia effect, are activated, partially counteracting the bradycardia effect.

### General limitations

A general limitation of our side-by-side characterization of recording electrode types for the peripheral nervous system is the varying degree of surgical trauma caused by the application of each electrode. The LIFE electrode insertion likely created the most trauma due to the large diameter insertion needle used to guide the electrode into the nerve, followed by the microneurography electrode, and lastly the cuff recording electrode, which created minimal additional trauma. Further, the open surgical preparation was not reflective of the minimally invasive clinical insertion typical with the microneurography electrode or the chronic healed-in recording state of an implanted cuff electrode. Another general limitation is that we did not fully optimize the design of each recording electrode type. However, many studies have already used computational and *in vivo* methods to optimize the design of each recording electrode type (Yoshida and Stein, 1999; Boretius et al., 2010; Sabetian et al., 2017; Jiman et al., 2020; Ottaviani et al., 2020; Blanz et al., 2022). We drew from these studies and prior experience to inform the electrode design for each electrode type tested. Despite these limitations, we believe the strong effect sizes of the primary outcomes give validity to the study’s conclusions.

## Conclusions

This publication is the first to report on a side-by-side comparison of electrodes for recording electrically evoked compound action potentials (ECAPs) and sensory evoked neural activity from the peripheral nervous system of a human-scaled large animal model. We characterized a recording cuff, microneurography electrodes, and longitudinal intrafascicular electrodes (LIFEs). We found that while the cuff records a significantly larger ECAP magnitude and has a lower noise floor than the microneurography electrode, the microneurography electrode may be the most sensitive recording electrode due to its proximity to the neural source. Further, the microneurography electrode consistently recorded A*β* ECAPs and was the only electrode to consistently record sensory evoked neural activity. We concluded that minimally invasive microneurography electrodes, already in routine human use, could be an important tool in the development and deployment of neuromodulation therapies to improve local on-target neural engagement.

## Acknowledgements

Dr. Jill Barnes for assistance with the microneurography technique and electrode selection. Dr. Rick Chappell for insights and input on statistical design and analysis. Maria LaLuzerne for swine study scheduling and animal care. Jonah Mudge for fabricating electrodes used in the pilot experiments. Carly Frieders, Ashlesha Deshmukh, Dr. Nicole Pelot, and Rex Chen for reviewing drafts of this manuscript. Members of the Wisconsin Institute for Translational Neuroengineering (WITNe) for feedback during group meetings. University of Wisconsin Translational Research Initiatives in Pathology laboratory (TRIP), supported by the UW Department of Pathology and Laboratory Medicine, UWCCC (P30 CA014520) and the Office of The Director-NIH (S10 OD023526) for use of its facilities and services to prepare the histology images.

This work was funded by the United States National Institute of Health (NIH) SPARC Program OT2OD025340.

## References

Ackerley, R., and Watkins, R. H. (2018). Microneurography as a tool to study the function of individual C-fiber afferents in humans: responses from nociceptors, thermoreceptors, and mechanoreceptors. Journal of Neurophysiology 120, 2834–2846. doi: 10.1152/jn.00109.2018.

Andreasen, L. N. S., and Struijk, J. J. (2002). Signal strength versus cuff length in nerve cuff electrode recordings. IEEE Trans. Biomed. Eng. 49, 1045–1050. doi: 10.1109/TBME.2002.800785.

Ardell, J. L., Nier, H., Hammer, M., Southerland, E. M., Ardell, C. L., Beaumont, E., et al. (2017). Defining the neural fulcrum for chronic vagus nerve stimulation: implications for integrated cardiac control: Vagus nerve stimulation neural fulcrum for cardiac control. J Physiol 595, 6887– 6903. doi: 10.1113/JP274678.

Barry, J. M. (2015). Axonal activity in vivo: technical considerations and implications for the exploration of neural circuits in freely moving animals. Front. Neurosci. 9. doi: 10.3389/fnins.2015.00153.

Blanz, S. L., Musselman, E. D., Settell, M. L., Knudsen, B. E., Nicolai, E. N., Trevathan, J. K., et al. (2022). Spatially selective stimulation of the pig vagus nerve to modulate target effect versus side effect. Bioengineering doi: 10.1101/2022.05.19.492726.

Boretius, T., Badia, J., Pascual-Font, A., Schuettler, M., Navarro, X., Yoshida, K., et al. (2010). A transverse intrafascicular multichannel electrode (TIME) to interface with the peripheral nerve. Biosensors and Bioelectronics 26, 62–69. doi: 10.1016/j.bios.2010.05.010.

Bovik, A. C., Acton, S. T. (2005). Handbook of Image and Video Processing (Second Edition). Academic Press, 99–108, ISBN 9780121197926. doi: 10.1016/B978-012119792-6/50070-X.

Buzsáki, G. (2004). Large-scale recording of neuronal ensembles. Nat Neurosci 7, 446–451. doi: 10.1038/nn1233.

Cakmak, Y. O., Apaydin, H., Kiziltan, G., Gündüz, A., Ozsoy, B., Olcer, S., et al. (2017). Rapid Alleviation of Parkinson’s Disease Symptoms via Electrostimulation of Intrinsic Auricular Muscle Zones. Front. Hum. Neurosci. 11, 338. doi: 10.3389/fnhum.2017.00338.

Castoro, M. A. (2011). Excitation properties of the right cervical vagus nerve in adult dogs. Experimental Neurology, 7.

Chakravarthy, K., Bink, H., and Dinsmoor, D. (2020). Sensing Evoked Compound Action Potentials from the Spinal Cord: Novel Preclinical and Clinical Considerations for the Pain Management Researcher and Clinician. JPR Volume 13, 3269–3279. doi: 10.2147/JPR.S289098.

Chakravarthy, K., FitzGerald, J., Will, A., Trutnau, K., Corey, R., Dinsmoor, D., et al. (2022). A Clinical Feasibility Study of Spinal Evoked Compound Action Potential Estimation Methods. Neuromodulation: Technology at the Neural Interface 25, 75–84. doi: 10.1111/ner.13510.

Chao, D., Mecca, C. M., Yu, G., Segel, I., Gold, M. S., Hogan, Q. H., et al. (2021). Dorsal root ganglion stimulation of injured sensory neurons in rats rapidly eliminates their spontaneous activity and relieves spontaneous pain. Pain 162, 2917–2932. doi: 10.1097/j.pain.0000000000002284.

Chen, H. (1994). Comparisons of Lognormal Population Means. Proceedings of the American Mathematical Society 121, 915. doi: 10.2307/2160293.

De Ferrari, G. M., Stolen, C., Tuinenburg, A. E., Wright, D. J., Brugada, J., Butter, C., et al. (2017). Long-term vagal stimulation for heart failure: Eighteen month results from the NEural Cardiac TherApy foR Heart Failure (NECTAR-HF) trial. International Journal of Cardiology 244, 229–234. doi: 10.1016/j.ijcard.2017.06.036.

Donaldson, N., Rieger, R., Schuettler, M., and Taylor, J. (2008). Noise and selectivity of velocity-selective multi-electrode nerve cuffs. Med Biol Eng Comput 46, 1005–1018. doi: 10.1007/s11517-008-0365-4.

Drebitz, E. (2020). A novel approach for removing micro-stimulation artifacts and reconstruction of broad-band neuronal signals. Journal of Neuroscience Methods, 11.

Duisit, J., Debluts, D., Behets, C., Gerdom, A., Vlassenbroek, A., Coche, E., et al. (2017). Porcine ear: A new model in large animals for the study of facial subunit allotransplantation. JPRAS Open 12, 47–58. doi: 10.1016/j.jpra.2017.01.004.

Erlanger, J., and Gasser, H. (1937). Electrical Signs of Nervous Activity (London: University of Pennsylvania Press)

Feucht, M., Ward, M. (2021). Non-invasive, Spatiotemporal Characterization of Muscle Activation Patterns from Vagus Nerve Stimulation in Human Subjects. Proceedings of IMPRS 4. doi: 10.18060/25920

Gaunt, R. A., Hokanson, J. A., and Weber, D. J. (2009). Microstimulation of primary afferent neurons in the L7 dorsal root ganglia using multielectrode arrays in anesthetized cats: thresholds and recruitment properties. J. Neural Eng. 6, 055009. doi: 10.1088/1741-2560/6/5/055009.

Getty, R., and Grossman, J. D. (1975). The Anatomy of the Domestic Animals, Vol. 2. W B Saunders Co. ISBN: 0721641075.

Gonzalez-Gonzalez, M. A., Bendale, G. S., Wang, K., Wallace, G. G., and Romero-Ortega, M. (2021). Platinized graphene fiber electrodes uncover direct spleen-vagus communication. Commun Biol 4, 1097. doi: 10.1038/s42003-021-02628-7.

Harding, G. W. (1991). A method for eliminating the stimulus artifact from digital recordings of the direct cortical response. Computers and Biomedical Research 24, 183–195. doi: 10.1016/0010-4809(91)90029-V.

He, S., Teagle, H. F. B., and Buchman, C. A. (2017). The Electrically Evoked Compound Action Potential: From Laboratory to Clinic. Front. Neurosci. 11, 339. doi: 10.3389/fnins.2017.00339.

Heusser, K., Tank, J., Brinkmann, J., Menne, J., Kaufeld, J., Linnenweber-Held, S., et al. (2016). Acute Response to Unilateral Unipolar Electrical Carotid Sinus Stimulation in Patients With Resistant Arterial Hypertension. Hypertension 67, 585–591. doi: 10.1161/HYPERTENSIONAHA.115.06486.

Humphrey, D., and Schmidt, E. (1990). Neurophysiological Techniques Applications to Neural Systems Neuromethods 15. Humana Totowa, ISBN: 978-1-4899-4117-6, doi: 10.1385/0896031853

Janko, M., and Trontelj, J. V. (1980). Transcutaneous Electrical Nerve Stimulation: A Microneurographic and Perceptual Study. Pain 9, 219–230. doi: 10.1016/0304-3959(80)90009-3

Jiman, A. A., Ratze, D. C., Welle, E. J., Patel, P. R., Richie, J. M., Bottorff, E. C., et al. (2020). Multi-channel intraneural vagus nerve recordings with a novel high-density carbon fiber microelectrode array. Sci Rep 10, 15501. doi: 10.1038/s41598-020-72512-7.

Johnson, R. L., and Wilson, C. G. (2018). A review of vagus nerve stimulation as a therapeutic intervention. JIR Volume 11, 203–213. doi: 10.2147/JIR.S163248.

Kaniusas, E., Kampusch, S., Tittgemeyer, M., Panetsos, F., Gines, R. F., Papa, M., et al. (2019). Current Directions in the Auricular Vagus Nerve Stimulation I – A Physiological Perspective. Front. Neurosci. 13, 854. doi: 10.3389/fnins.2019.00854.

Khodagholy, D., Gelinas, J. N., Thesen, T., Doyle, W., Devinsky, O., Malliaras, G. G., et al. (2015). NeuroGrid: recording action potentials from the surface of the brain. Nat Neurosci 18, 310–315. doi: 10.1038/nn.3905.

Kodandaramaiah, S. B., Franzesi, G. T., Chow, B. Y., Boyden, E. S., and Forest, C. R. (2012). Automated whole-cell patch-clamp electrophysiology of neurons in vivo. Nat Methods 9, 585–587. doi: 10.1038/nmeth.1993.

Koh, R. G. L., Zariffa, J., Jabban, L., Yen, S.-C., Donaldson, N., and Metcalfe, B. W. (2022). Tutorial: A guide to techniques for analysing recordings from the peripheral nervous system. J. Neural Eng. doi: 10.1088/1741-2552/ac7d74.

Koppaka, S., Hess-Dunning, A., and Tyler, D. J. (2022). Biomechanical characterization of isolated epineurial and perineurial membranes of rabbit sciatic nerve. Journal of Biomechanics 136, 111058. doi: 10.1016/j.jbiomech.2022.111058.

Lefkowitz, T., Hazani, R., Chowdhry, S., Elston, J., Yaremchuk, M. J., and Wilhelmi, B. J. (2013). Anatomical Landmarks to Avoid Injury to the Great Auricular Nerve During Rhytidectomy. Aesthetic Surgery Journal 33, 19–23. doi: 10.1177/1090820X12469625.

Lundberg, K. H. (2002). Noise Sources in Bulk CMOS. MIT web accessed at https://web.mit.edu/klund/www/papers/UNP_noise.pdf

Macefield, V. G. (2021). Recording and quantifying sympathetic outflow to muscle and skin in humans: methods, caveats and challenges. Clin Auton Res 31, 59–75. doi: 10.1007/s10286-020-00700-6.

Manzano, G. M., Giuliano, L. M. P., and Nóbrega, J. A. M. (2008). A brief historical note on the classification of nerve fibers. Arq. Neuro-Psiquiatr. 66, 117–119. doi: 10.1590/S0004-282X2008000100033.

Meier, K., Qerama, E., Ettrup, K. S., Glud, A. N., Alstrup, A. K. O., and Sørensen, J. C. H. (2018). Segmental innervation of the Göttingen minipig hind body. An electrophysiological study. J. Anat. 233, 411–420. doi: 10.1111/joa.12865.

Mekhail, N., Levy, R. M., Deer, T. R., Kapural, L., Li, S., Amirdelfan, K., et al. (2022). Durability of Clinical and Quality-of-Life Outcomes of Closed-Loop Spinal Cord Stimulation for Chronic Back and Leg Pain: A Secondary Analysis of the Evoke Randomized Clinical Trial. JAMA Neurol 79, 251. doi: 10.1001/jamaneurol.2021.4998.

Metcalfe, B. W., Nielsen, T. N., Donaldson, N. de N., Hunter, A. J., and Taylor, J. T. (2018). First demonstration of velocity selective recording from the pig vagus using a nerve cuff shows respiration afferents. Biomed. Eng. Lett. 8, 127–136. doi: 10.1007/s13534-017-0054-z.

Mitra, S., Mitra, M., and Halder, B. (2018). Automatic feature extraction of ECG signal based on adaptive window dependent differential histogram approach and validation with CSE database. IJCSYSE 4, 146. doi: 10.1504/IJCSYSE.2018.10012644.

Molecular Devices. (2006). The Axon CNS Guide. 235–263, Part Number 2500-102 Rev B 200

Moncrief, K., and Kaufman, S. (2006). Splenic baroreceptors control splenic afferent nerve activity. American Journal of Physiology-Regulatory, Integrative and Comparative Physiology 290, R352– R356. doi: 10.1152/ajpregu.00489.2005.

Nicolai, E. N., Settell, M. L., Knudsen, B. E., McConico, A. L., Gosink, B. A., Trevathan, J. K., et al. (2020). Sources of off-target effects of vagus nerve stimulation using the helical clinical lead in domestic pigs. J. Neural Eng. 17, 046017. doi: 10.1088/1741-2552/ab9db8.

Nonis, R., D’Ostilio, K., Schoenen, J., and Magis, D. (2017). Evidence of activation of vagal afferents by non-invasive vagus nerve stimulation: An electrophysiological study in healthy volunteers. Cephalalgia 37, 1285–1293. doi: 10.1177/0333102417717470.

Obot, I. B., Onyeachu, I. B., Zeino, A., and Umoren, S. A. (2019). Electrochemical noise (EN) technique: review of recent practical applications to corrosion electrochemistry research. Journal of Adhesion Science and Technology 33, 1453–1496. doi: 10.1080/01694243.2019.1587224.

Ottaviani, M. M., Wright, L., Dawood, T., and Macefield, V. G. (2020). In vivo recordings from the human vagus nerve using ultrasound-guided microneurography. J Physiol, JP280077. doi: 10.1113/JP280077.

Park, J., Choi, W.-M., Kim, K., Jeong, W.-I., Seo, J.-B., and Park, I. (2018). Biopsy Needle Integrated with Electrical Impedance Sensing Microelectrode Array towards Real-time Needle Guidance and Tissue Discrimination. Sci Rep 8, 264. doi: 10.1038/s41598-017-18360-4.

Parker, J. L., Karantonis, D. M., Single, P. S., Obradovic, M., and Cousins, M. J. (2012). Compound action potentials recorded in the human spinal cord during neurostimulation for pain relief. Pain 153, 593–601. doi: 10.1016/j.pain.2011.11.023.

Parker, J. L., Obradovic, M., Hesam Shariati, N., Gorman, R. B., Karantonis, D. M., Single, P. S., et al. (2020). Evoked Compound Action Potentials Reveal Spinal Cord Dorsal Column Neuroanatomy. Neuromodulation: Technology at the Neural Interface 23, 82–95. doi: 10.1111/ner.12968.

Pelot, N. A., Behrend, C. E., and Grill, W. M. (2019). On the parameters used in finite element modeling of compound peripheral nerves. J. Neural Eng. 16, 016007. doi: 10.1088/1741-2552/aaeb0c.

Pelot, N. A., Goldhagen, G. B., Cariello, J. E., Musselman, E. D., Clissold, K. A., Ezzell, J. A., et al. (2020). Quantified Morphology of the Cervical and Subdiaphragmatic Vagus Nerves of Human, Pig, and Rat. Front. Neurosci. 14, 601479. doi: 10.3389/fnins.2020.601479.

Plonsey, R., and Barr, R. C. (1995). Electric field stimulation of excitable tissue. IEEE Trans. Biomed. Eng. 42, 329–336. doi: 10.1109/10.376126.

Qing, K. Y., Wasilczuk, K. M., Ward, M. P., Phillips, E. H., Vlachos, P. P., Goergen, C. J., et al. (2018). B fibers are the best predictors of cardiac activity during Vagus nerve stimulation: Qing, vagal B fiber activation and cardiac effects. Bioelectron Med 4, 5. doi: 10.1186/s42234-018-0005-8.

Rawlins, J. C. (2000). “AC and the Sine Wave,” in Basic AC Circuits (Elsevier), 29–63. doi: 10.1016/B978-075067173-6/50003-1.

Ringer, S. K., Spielmann, N., Weiss, M., and Mauch, J. Y. (2016). Fentanyl bolus induces muscle tremors in sevoflurane-anaesthetized piglets. Lab Anim 50, 312–314. doi: 10.1177/0023677215623896.

Robinson, D. A. (1968). The electrical properties of metal microelectrodes. Proc. IEEE 56, 1065–1071. doi: 10.1109/PROC.1968.6458.

Sabbah, H. N., Ilsar, I., Zaretsky, A., Rastogi, S., Wang, M., and Gupta, R. C. (2011). Vagus nerve stimulation in experimental heart failure. Heart Fail Rev 16, 171–178. doi: 10.1007/s10741-010-9209-z.

Sabetian, P., Popovic, M. R., and Yoo, P. B. (2017). Optimizing the design of bipolar nerve cuff electrodes for improved recording of peripheral nerve activity. J. Neural Eng. 14, 036015. doi: 10.1088/1741-2552/aa6407.

Scipy. (2022). scipy.signal.filtfilt. Scipy v1.9.1 Manual accessed at https://docs.scipy.org/doc/scipy/reference/generated/scipy.signal.filtfilt.html

Settell, M. L., Pelot, N. A., Knudsen, B. E., Dingle, A. M., McConico, A. L., Nicolai, E. N., et al. (2020). Functional vagotopy in the cervical vagus nerve of the domestic pig: implications for the study of vagus nerve stimulation. J. Neural Eng. 17, 026022. doi: 10.1088/1741-2552/ab7ad4.

Sharma, V. S., Stephens, R. E., Wright, B. W., and Surek, C. C. (2017). What Is the Lobular Branch of the Great Auricular Nerve? Anatomical Description and Significance in Rhytidectomy: Plastic and Reconstructive Surgery 139, 371e–378e. doi: 10.1097/PRS.0000000000002980.

Silverman, H. A., Stiegler, A., Tsaava, T., Newman, J., Steinberg, B. E., Masi, E. B., et al. (2018). Standardization of methods to record Vagus nerve activity in mice. Bioelectron Med 4, 3. doi: 10.1186/s42234-018-0002-y.

Simakov, A. B., and Webster, J. G. (2010). Motion Artifact From Electrodes and Cables. Iranian Journal of Electrical and Computer Engineering 9, 139–143.

Stumpp, L., Smets, H., Vespa, S., Cury, J., Doguet, P., Delbeke, J., et al. (2020). Recording of spontaneous vagus nerve activity during Pentylenetetrazol-induced seizures in rats. Journal of Neuroscience Methods 343, 108832. doi: 10.1016/j.jneumeth.2020.108832.

Tucker-Davis Technologies, Inc. (TDT). (2022). SIM Subject Interface Module Hardware Reference, Page 23

Ulrich, G., Schlosser, W., and Juckel, G. (2016). The impact of Arterial Pulse Impedance Artifact (APIA) on test-retest reliability of quantitative EEG. Neuropsychiatr Electrophysiol 2, 5. doi: 10.1186/s40810-016-0019-y.

Verma, N., Graham, R. D., Mudge, J., Trevathan, J. K., Franke, M., Shoffstall, A. J., et al. (2021a). Augmented Transcutaneous Stimulation Using an Injectable Electrode: A Computational Study. Front. Bioeng. Biotechnol. 9, 796042. doi: 10.3389/fbioe.2021.796042.

Verma, N., Mudge, J. D., Kasole, M., Chen, R. C., Blanz, S. L., Trevathan, J. K., et al. (2021b). Auricular Vagus Neuromodulation—A Systematic Review on Quality of Evidence and Clinical Effects. Front. Neurosci. 15, 664740. doi: 10.3389/fnins.2021.664740.

Winestone, J. S., Zaidel, A., Bergman, H., and Israel, Z. (2012). The use of macroelectrodes in recording cellular spiking activity. Journal of Neuroscience Methods 206, 34–39. doi: 10.1016/j.jneumeth.2012.02.002.

Yang, H.-M., Kim, H.-J., and Hu, K.-S. (2015). Anatomic and histological study of great auricular nerve and its clinical implication. Journal of Plastic, Reconstructive & Aesthetic Surgery 68, 230–236. doi: 10.1016/j.bjps.2014.10.030.

Yang, Z., Zhao, Q., Keefer, E., and Liu, W. (2009). Noise Characterization, Modeling, and Reduction for In Vivo Neural Recording. NIPS Advances in Neural Information Processing Systems 22. ISBN: 9781615679119

Yasar, N. (2021). Causes of noise in electrophysiological recordings. Plexon blog accessed at https://plexon.com/blog-post/causes-of-noise-in-electrophysiological-recordings/

Yoo, P. B., Lubock, N. B., Hincapie, J. G., Ruble, S. B., Hamann, J. J., and Grill, W. M. (2013). High-resolution measurement of electrically-evoked vagus nerve activity in the anesthetized dog. J. Neural Eng. 10, 026003. doi: 10.1088/1741-2560/10/2/026003.

Yoshida, K., and Stein, R. B. (1999). Characterization of signals and noise rejection with bipolar longitudinal intrafascicular electrodes. IEEE Trans. Biomed. Eng. 46, 226–234. doi: 10.1109/10.740885.

