## Supplementary materials for "Microneurography as a Minimally Invasive Method to Assess Target Engagement During Neuromodulation"

**Supplementary Material 1 – Close-up photographs of recording electrodes**

**Supplementary Material 2 – Histology of great auricular nerve**

**Supplementary Material 3 – Great auricular nerve surgical pocket**

**Supplementary Material 4 – Facial nerve anatomy**

**Supplementary Material 5 – Great auricular nerve anatomy**

**Supplementary Material 6 – Non-functional electrodes**

**Supplementary Material 7 – Detrending of ECAPs from stimulation artifact**

**Supplementary Material 8 – Primary noise contribution in time series from low-frequency noise (e.g., cardiac related artifacts)**

**Supplementary Material 9 – Verifying authenticity of B-fiber recordings using signal propagation delay**

**Supplementary Material 10 – Data from all subjects**

**Supplementary Material 11 – Spontaneous activity recordings from cervical vagus nerve (cVN)**

**Supplementary Material 1 – Close-up photographs of recording electrodes**

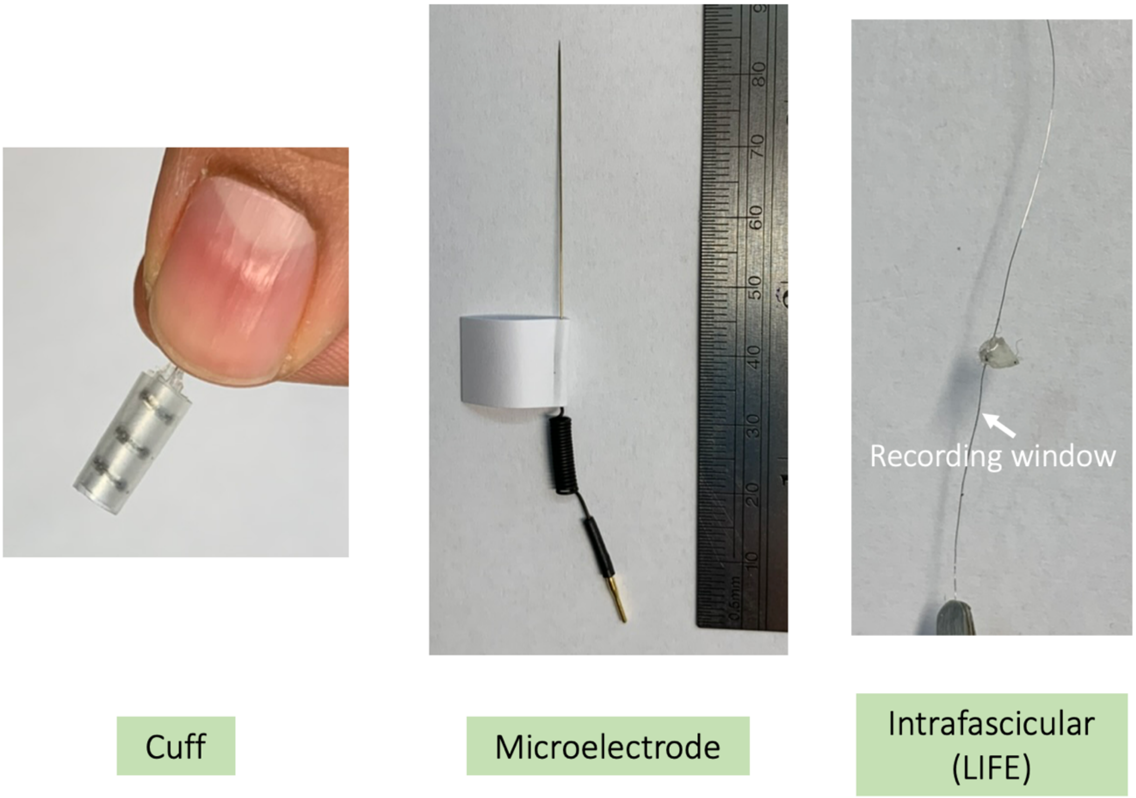


**Figure S1-1:** Close up photograph of three recording electrodes characterized in this study.

**Supplementary Material 2 – Histology of great auricular nerve**

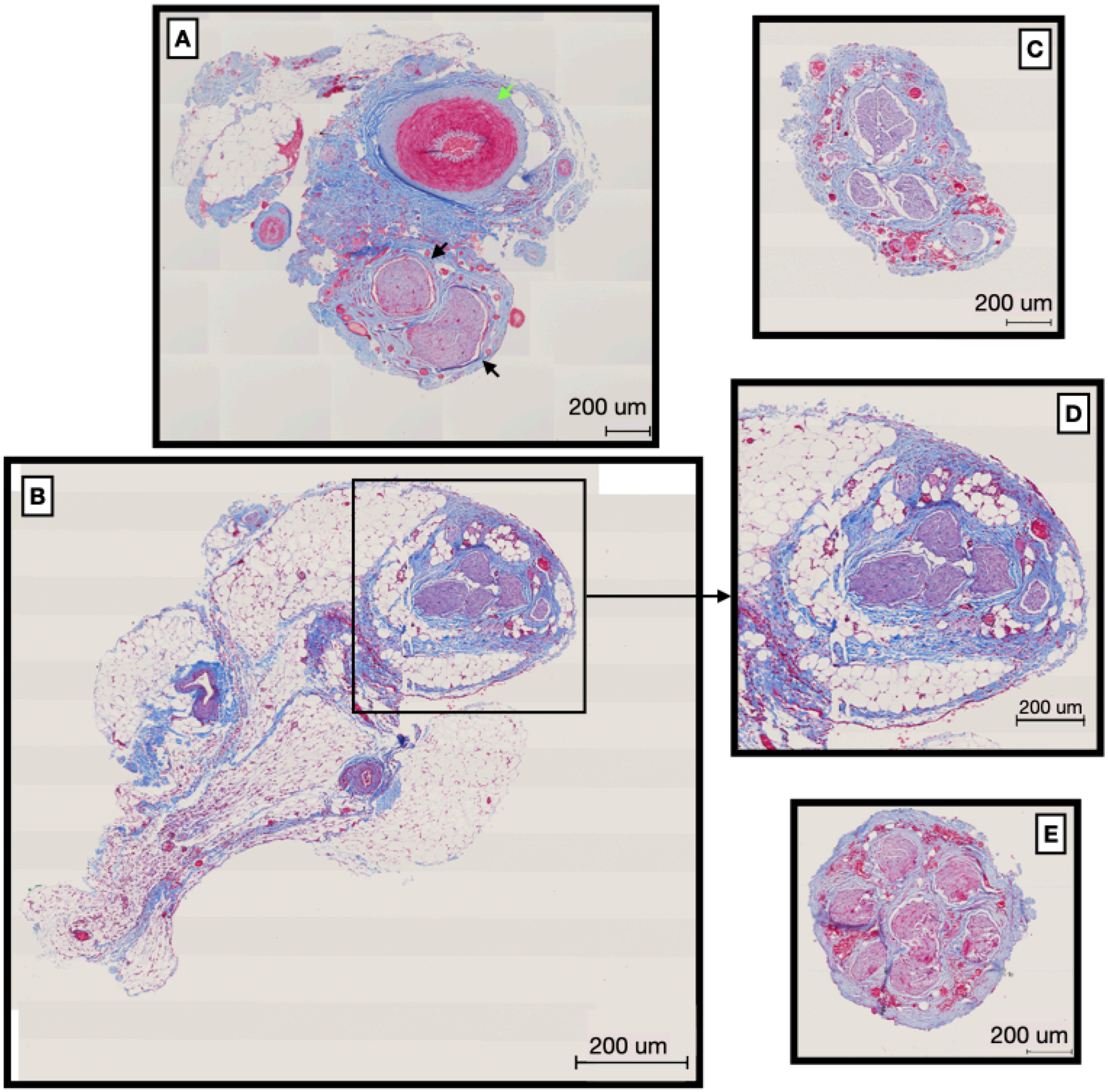


**Figure S2-1:** Representative histology of great auricular nerve cross sections (5 um) for four subjects. **(A)** Contains both the fascicles of the great auricular nerve (black arrow heads) and the associated artery (green arrowhead) **(D)** Zoomed region of **(B)**, showing fascicles of the great auricular nerve.

**Supplementary Material 3 – Great auricular nerve surgical pocket**

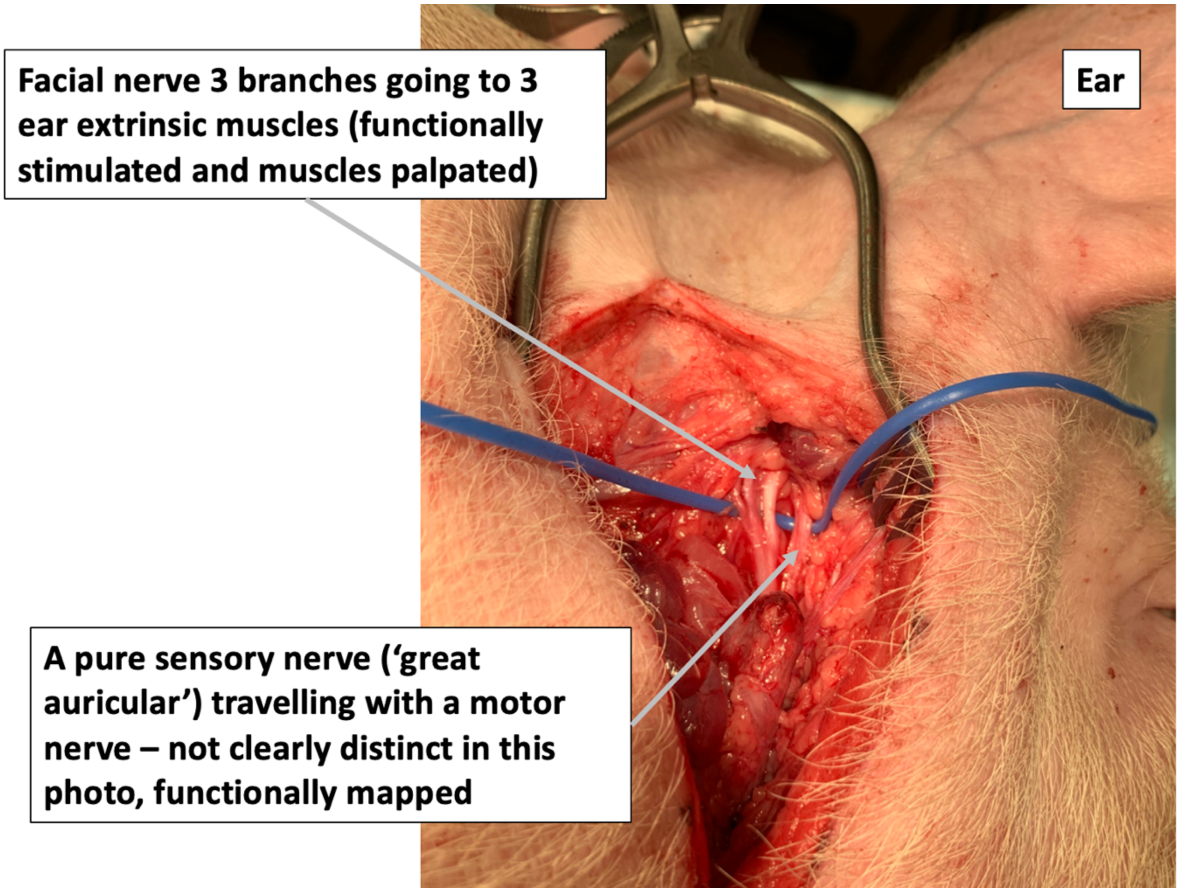


**Figure S3-1:** Surgical pocket to access great auricular nerve and surround anatomy. Functional mapping allowed differentiation between motor nerves (facial) and the sensory nerve (great auricular)

**Supplementary Material 4 – Facial nerve anatomy**

**Facial nerve (pig):**

1. caudal auricular nerve
2. internal auricular nerve
3. auriculopalpebral nerve
4. dorsal buccal branch
5. ventral buccal branch
6. stylohyoid branch
7. cervical branch

Note in human literature, ‘anterior’ and ‘posterior’ are used while in animal literature ‘rostral’ and ‘caudal’ are used respectively.

**Caudal/posterior auricular nerve**:
Initiates near stylomastoid foramen courses dorsal bifurcating into caudal and rostral branches. Both caudal and rostral branches course in association with caudal auricular artery and internal auricular nerve. Caudal branch is smaller of the two and innervates cervicoauricularis profundus muscle, with smaller branches innervating transverse and oblique auricular muscles as well as parotidoauricularis muscle. Innervates both intrinsic and extrinsic muscles of the outer ear.

**Rostral/anterior auricular branch:**
Continues a dorsal course and passes between retroauricular fat pad and ventral surface of the cervicoauricularis profundus muscle. In this area the rostral branch gives off smaller branches to styloauricularis muscle. These branches penetrate the auricular cartilage at helicine fissure and innervate the helicine muscle, terminating in the styloauricularis m.

**Internal auricular branch:**
Originates from dorsal margin of facial nerve and courses dorsal.
Divides into two to three branches before entering auricular cartilage and ramifying the skin of the inner surface of the auricular cartilage.

**Auriculopalpebral nerve:**
Courses dorsal communicating with auriculotemporal nerve then dividing into rostral auricular branches which innervate the auriculares rostrales muscles and a zygomatic branch to the frontoscutularis, levator anguliocusli medialis and orbicularis oculi muscles.

*Sisson and Grossman’s “The Anatomy of the Domestic Animals”, 1975.

**Supplementary Material 5 – Great auricular nerve anatomy**

The great auricular nerve (GAN) courses dorsally after emerging from the sternomastoid muscle. Traveling below the platysma muscle and coursing in association with the external jugular vein to the parotid margin. At this point the GAN begins to divide into its terminal branches. The GAN bifurcates into two primary branches with each of the primary branches dividing further.

The first division is the anterior branch. This branch courses rostrally over the parotid gland bifurcating into superficial and deep branches. The superficial branch innervates the skin over the parotid gland on the face. The deep branch penetrates the SMAS and parotid fascia and communicates with the facial nerve within the parotid gland (Yang et.al JPRAS 2015).

The posterior branch continues dorsal towards the posterior base of the ear and in association with the lateral auricular vein. It continues coursing around the posterior margin of the ear branching out to innervate the skin over the mastoid process. Another branch enters the cartilage at the base of the ear and supplies the anterior lower third of the ear. In humans this branch has been identified and referred to as the lobular branch (Sharma et al 2016). In pigs the lobular or lower third of the ear does not have a fatty lobule, the lobular equivalent of the pig would include the exterior skin around the tragus, antitragus, and intertragic notch areas of the ear. The posterior branch then continues along the posterior/caudal margin of the ear to innervate the posterior surface/skin of the auricle.

Some studies have suggested that this “lobular” branch be identified as a separate branch that supplies sensory innervation to the lower third of the ear. There are several studies that suggest that the GAN branches have four to five main configurations, we however did not dissect the entire configuration from each pig but did note that locations for the nerve configurations varied between animals (Sharma et al ASPS 2016; Lefkowitz et al., Aesthetic Surgery Journal, 2013; Yang et al. JPRAS 2015).

We found very little information on peripheral nerve anatomy or function in the pig model. Furthermore, the information can be misleading and inconsistent compared with surgical observations. We did not conduct a complete anatomical study for our purposes. However, future studies are warranted if the GAN in pig is to become a standard model.

**Supplementary Material 6 – Non-functional electrodes**

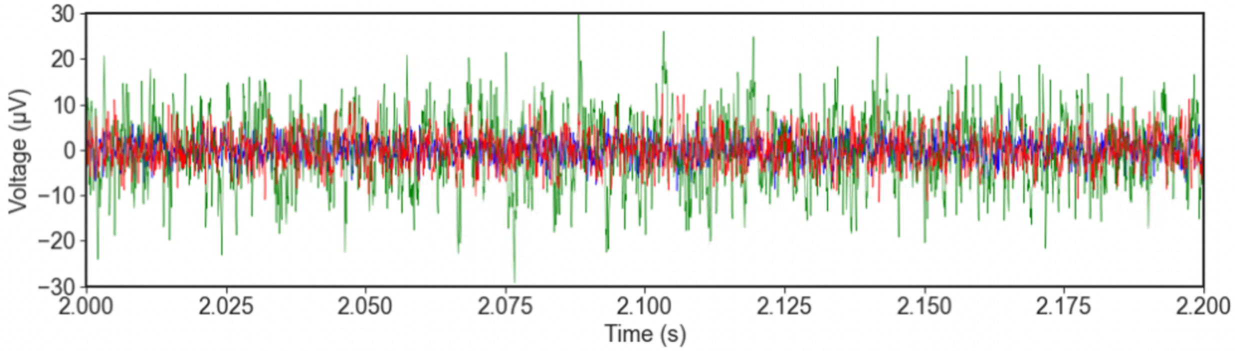


**Figure S6-1: Subject 1 vagus micro 2** was a non-functional electrode classified by the high noise floor (green trace) compared to the blue and red traces (figure below) and no ECAP signal even at 5 mA of stimulation

**Subject 1 great auricular micro 2** was a non-functional electrode classified by the high noise floor (figure in Supplementary Material 10 – Data from all subjects).

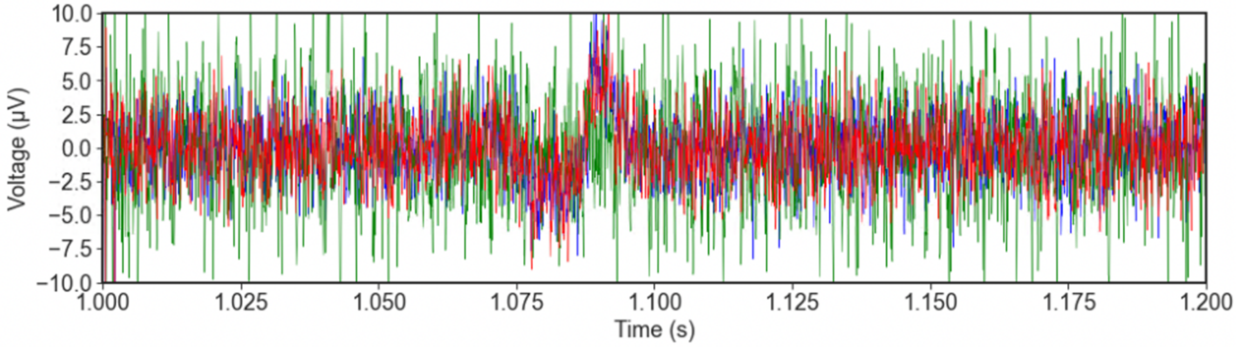


**Figure S6-2: Subject 4 vagus cuff 2** was a non-functional electrode classified by the high noise floor (green trace) compared to the blue and red traces (figure below) and noted as physically disconnected at the end of the recording.

**Supplementary Material 7 – Detrending of ECAPs from stimulation artifact**

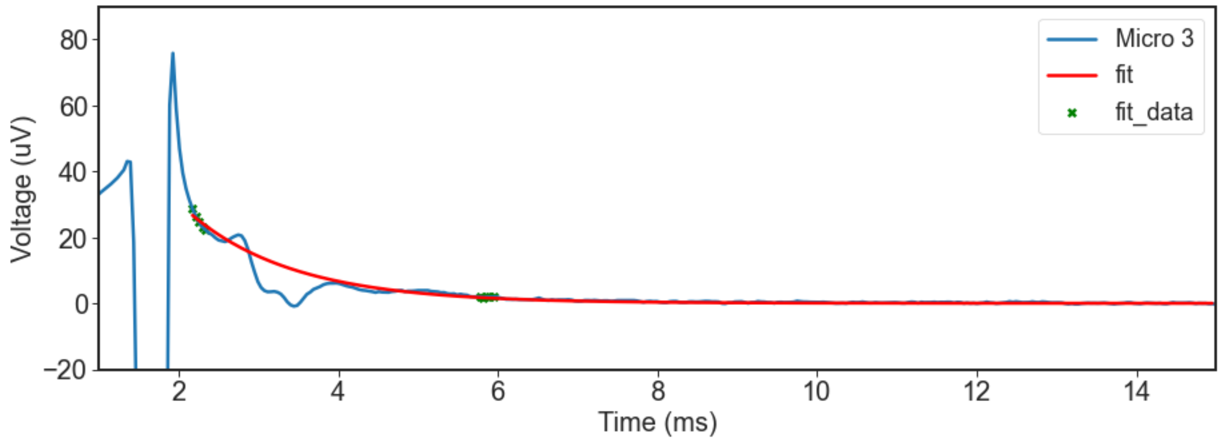


**Figure S7-1:** Only ECAPs from Subject 6 microelectrode recordings during the vagus dose-response amplitude sweep were contaminated by stimulation artifact. This is a representative fit of an exponential decay to the stimulation artifact in the ECAP trace.

**Supplementary Material 8 – Primary noise contribution in time series from low-frequency noise (e.g., cardiac related artifacts)**

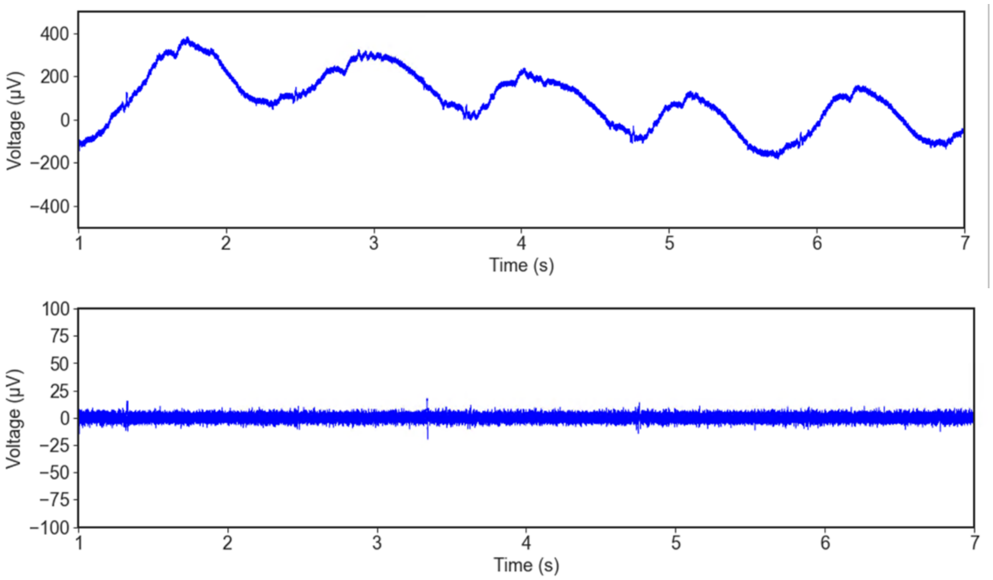


**Figure S8-1:** Microelectrode traces from subject 1 before (top) and after (bottom) filtering. High RMS noise in unfiltered microelectrode time series is coming largely from low frequencies.

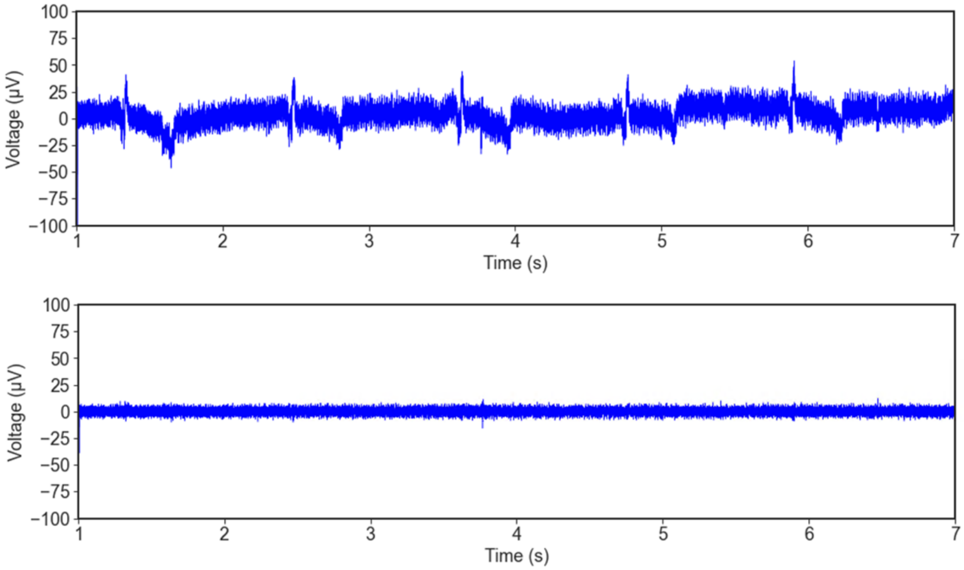


**Figure S8-2:** (top) An example of cardiac related artifact appearing in electrophysiology recordings, which may be mistaken for a neural signal. The cardiac related effect could be cardioballistic in nature or from the electrical activity of the heart, depending on the setup of the reference and recording electrode. (bottom) Removed by high pass filtering (fc = 100 Hz). See Fig. 5B in main paper for additional artifact examples. Cuff traces from subject 1.

**Supplementary Material 9 – Verifying authenticity of B-fiber recordings using signal propagation delay**

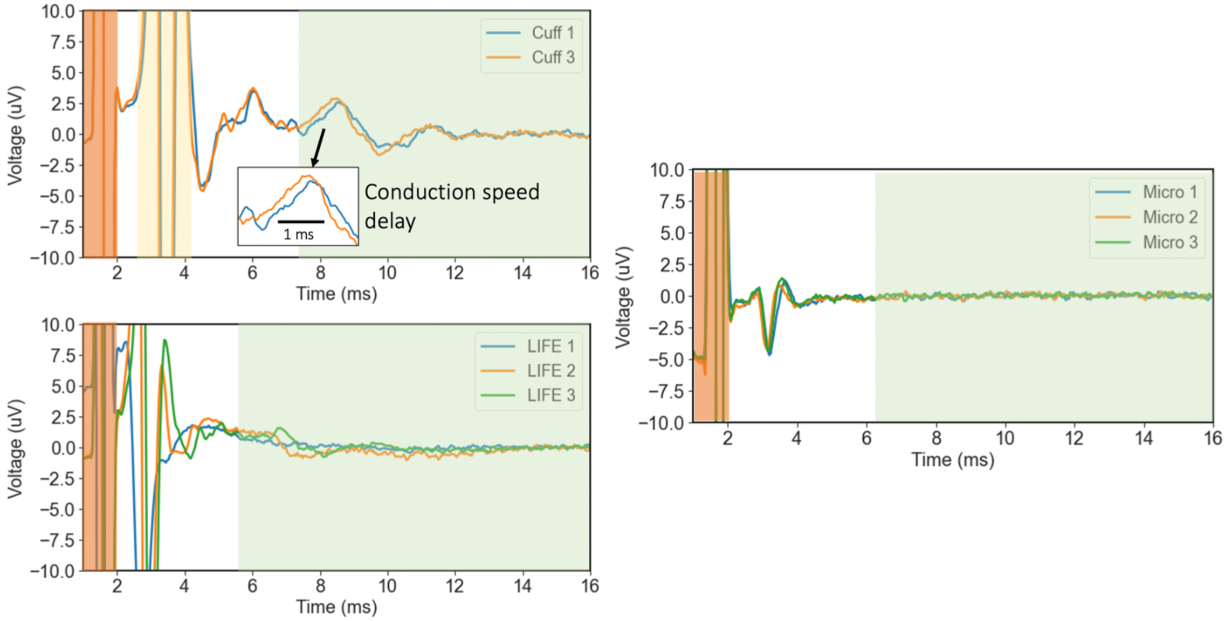


**Figure S9-1:** ECAP during 10 mA of stimulation on the vagus nerve in subject 4. (top left) with recording cuff (bottom left) with intrafascicular electrodes and (right) with microneurography microelectrode. Stimulation artifact shaded in orange, A$\beta$-fiber ECAP shaded in yellow, and B-fiber ECAP region shaded in green. Authenticity of B-fiber ECAP is suggested by signal propagation delay in cuff and intrafascicular electrode recording but not in microneurography microelectrode recording.

### Supplementary Material 10 – Data from all subjects

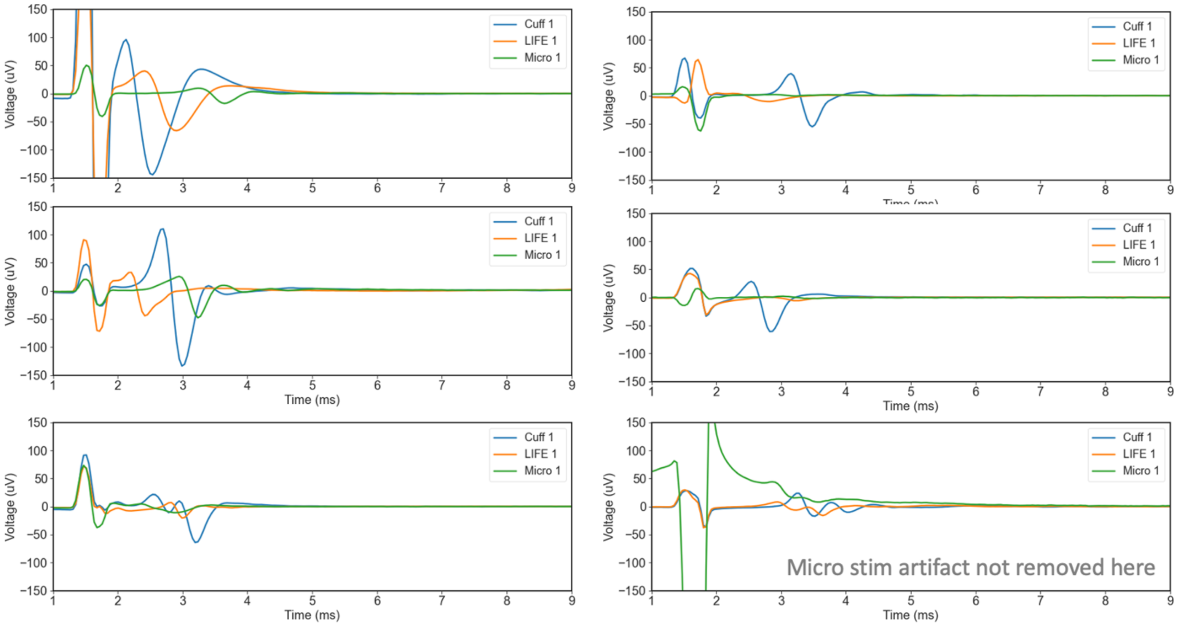


**Figure S10-1:** A$\beta$ ECAP at 1.5 mA of stimulation on cervical vagus nerve from all subjects. (top-left) subject 1 to (bottom-left) subject 3 and (top-right) subject 4 to (bottom-right) subject 6. Cuff consistently records the largest ECAP. Apparent triphasic morphology in subject 1 cuff may be due to stimulation artifact recovery.

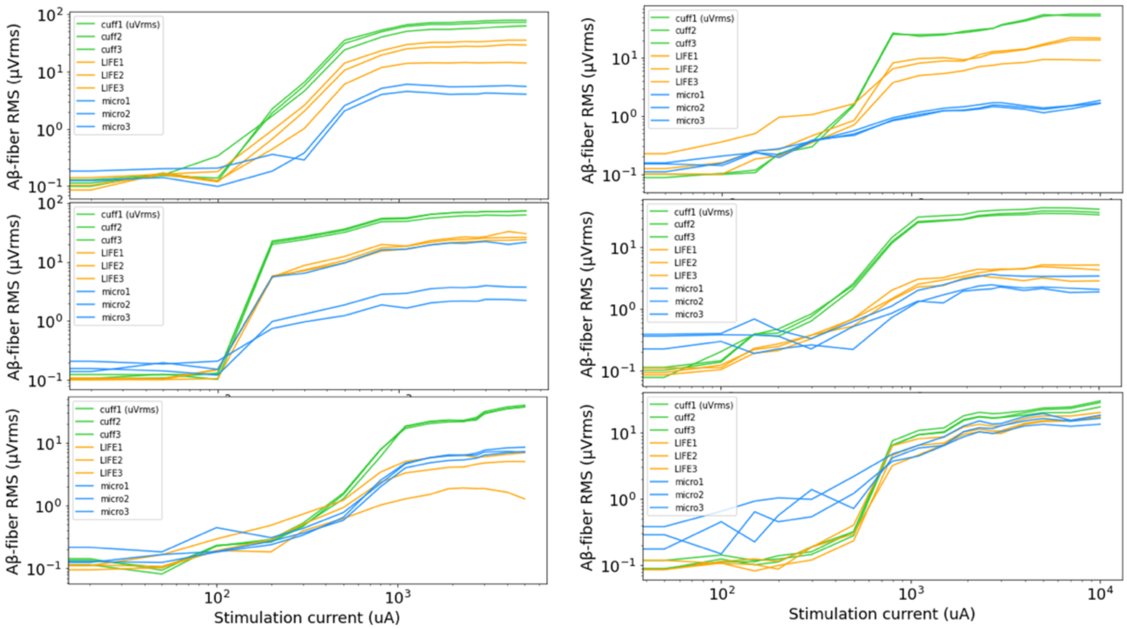


**Figure S10-2:** A$\beta$-fiber dose-response curves during vagus nerve stimulation from all subjects. (top-left) subject 1 to (bottom-left) subject 3 and (top-right) subject 4 to (bottom-right) subject 6.

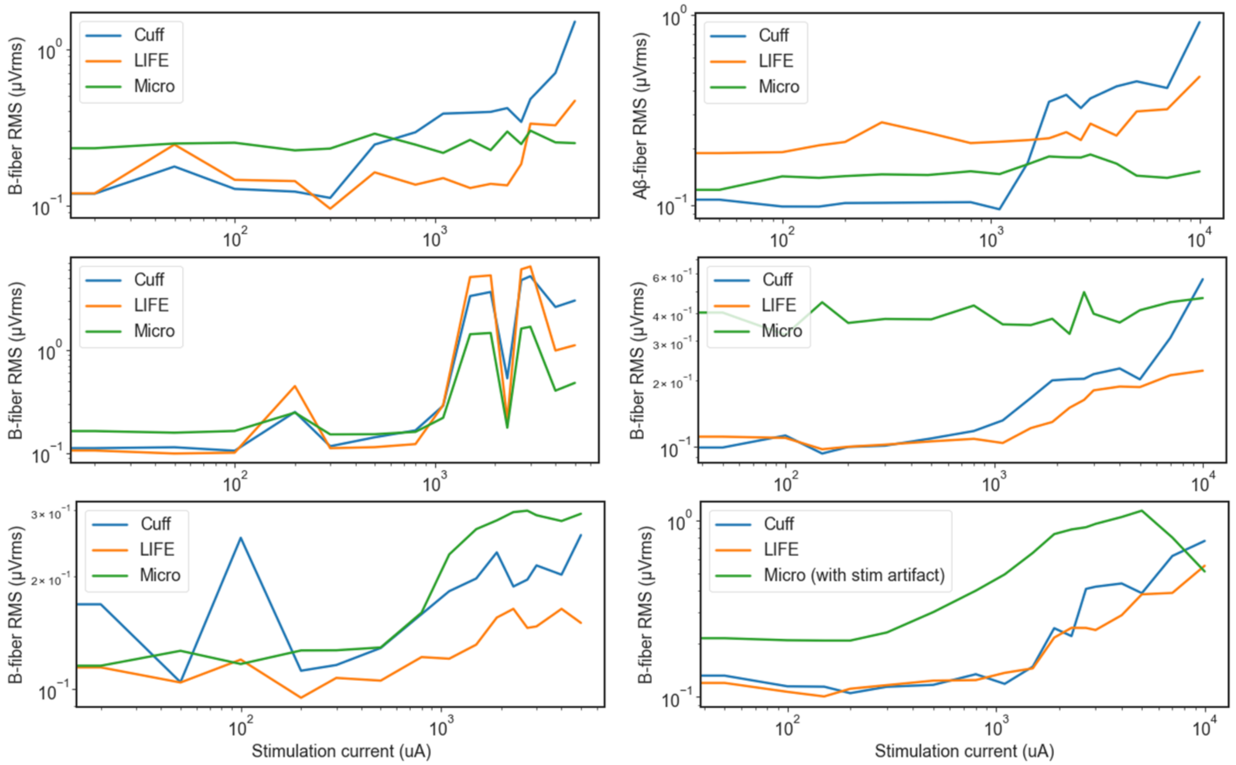


**Figure S10-3:** B-fiber dose-response curves during vagus nerve stimulation from all subjects. (top-left) subject 1 to (bottom-left) subject 3 and (top-right) subject 4 to (bottom-right) subject 6. An insufficient dose of Vecuronium (muscle blocker) was used in subjects 1-3 so EMG contamination of the B-fiber dose-response curves is likely. Not subject 6 microneurography microelectrode recordings are contaminated by stimulation artifact.

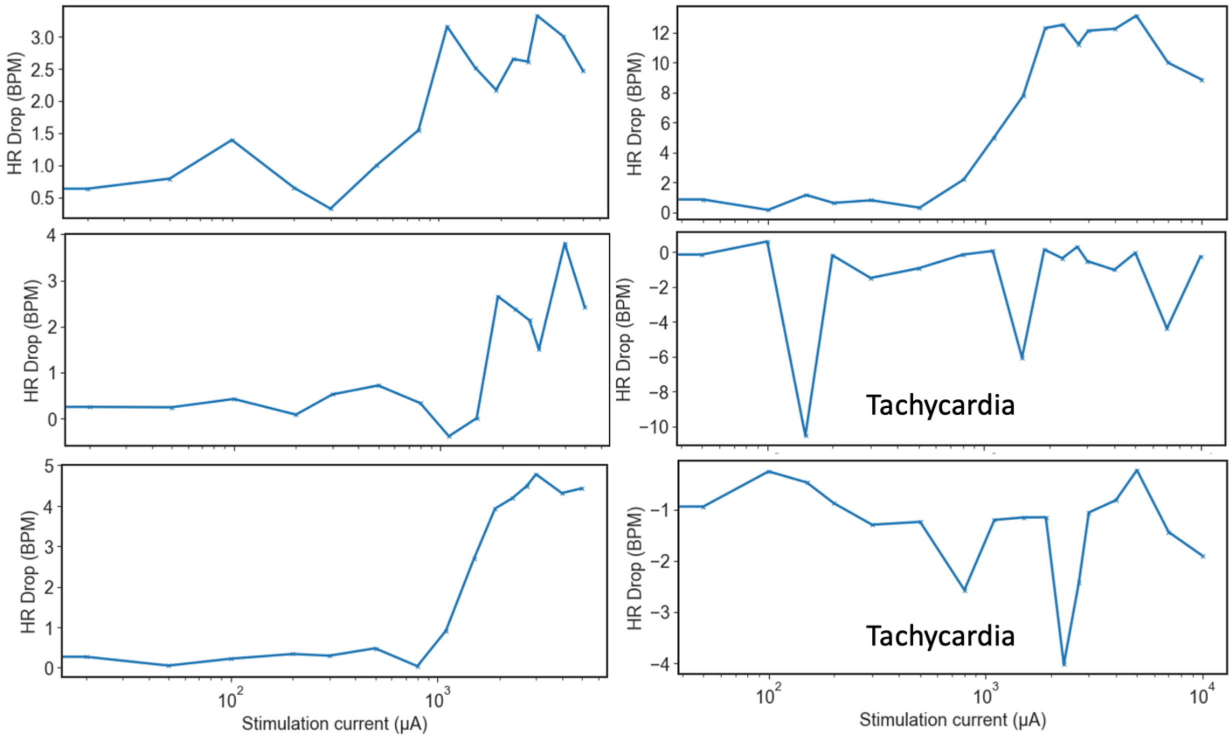


**Figure S10-4:** Stimulated evoked heart rate change dose-response curves during vagus nerve stimulation (VNS) from all subjects. (top-left) subject 1 to (bottom-left) subject 3 and (top-right) subject 4 to (bottom-right) subject 6. Subjects 5 and 6 are showing a tachycardia response instead of the canonical bradycardia response to VNS. Note that subjects 1-3 had the vagus experiment performed in the afternoon and were under greater doses of anesthesia compared to subjects 4-6 that had the vagus experiment performed in the morning and were under lower doses of anesthesia.

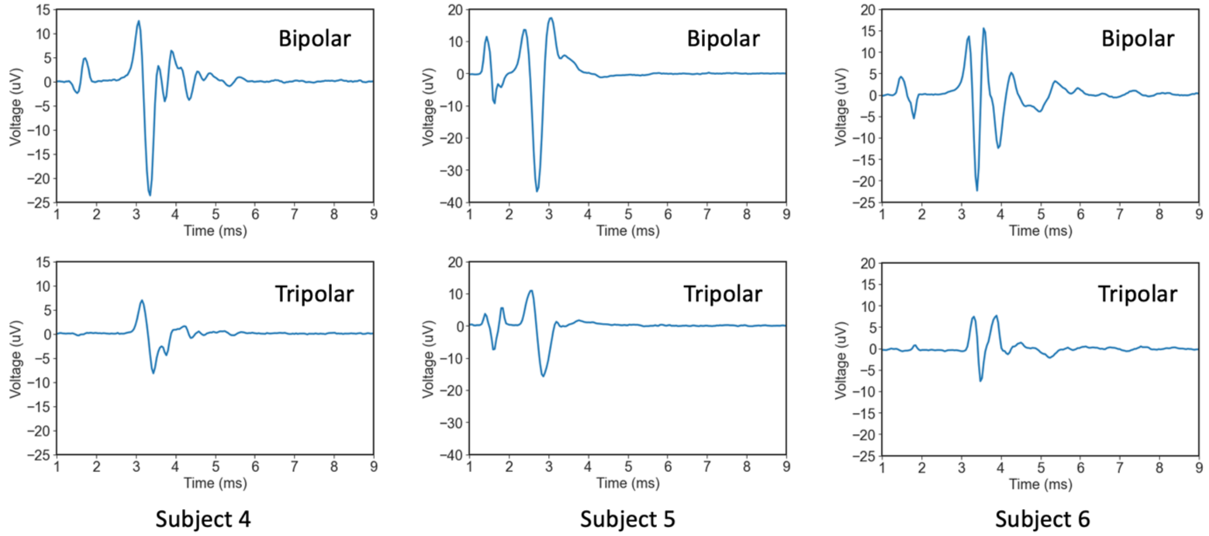


**Figure S10-5:** Tripolar vs. bipolar reference cuff recordings from subjects 4 (left), 5 (center), and 6 (right). Stimulation artifact is consistently smaller in tripolar. ECAP magnitude is consistently smaller in tripolar by factor of ~2.2x compared to bipolar reference.

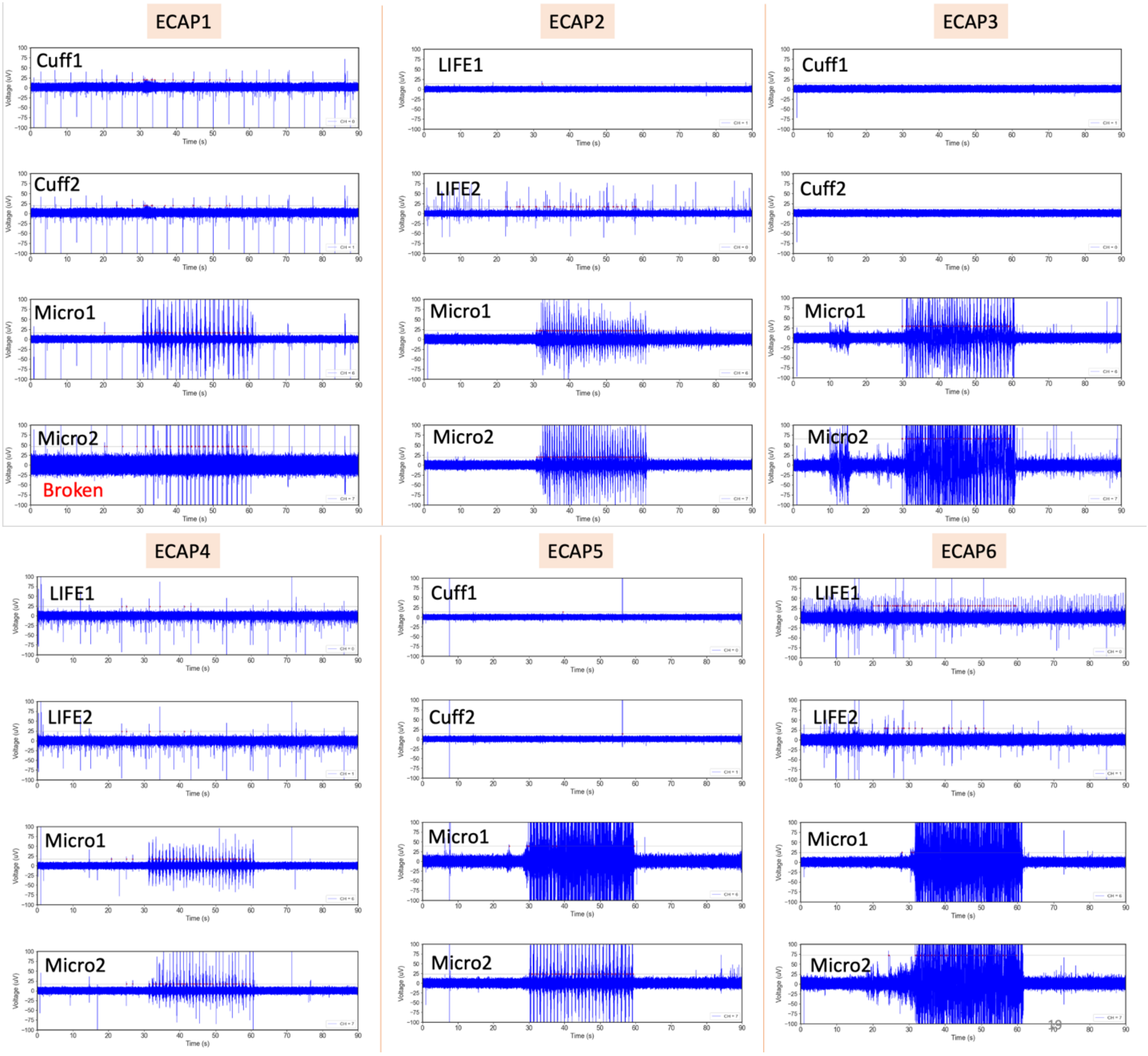


**Figure S10-6**: Spike count from sensory stroking evoked naturally occurring activity on all subjects. (top-left) subject 1 to (bottom-right) subject 6. 90 seconds of recording shown: first 30 seconds are quiescent with no stroking, 30-60s are on-target stroking at the region of the auricle innervated by the great auricular nerve, and 60-90s are off-target stroking at a region of the auricle not innervated by the great auricular nerve.

**Supplementary Material 11 – Spontaneous activity recordings from cervical vagus nerve (cVN)**

None of the electrode recordings showed clear and repeatable spontaneously occurring neural activity in the cVN. Signals that initially appeared neural were likely motion artifacts as they persisted more than 20 minutes after the nerve was transected cranial and caudal to the recording electrodes. Further, the artifacts were more pronounced in subjects 1-3, where the subjects experienced more tremor as insufficient muscle paralytic, vecuronium, was administered, compared to subjects 4-6.

Representative recordings from 1 cuff contact (channel 0), 1 LIFE electrode (channel 5), and 1 microneurography electrode (channel 8) per subject:

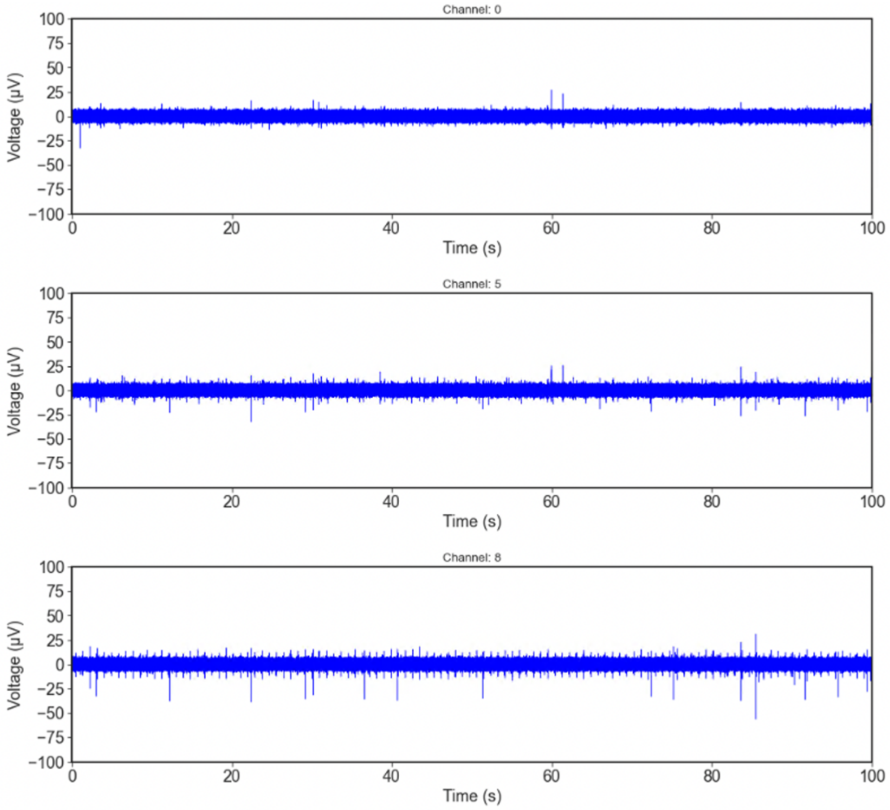


**Figure S11-1:** Subject 1 cVN spontaneous activity recording.

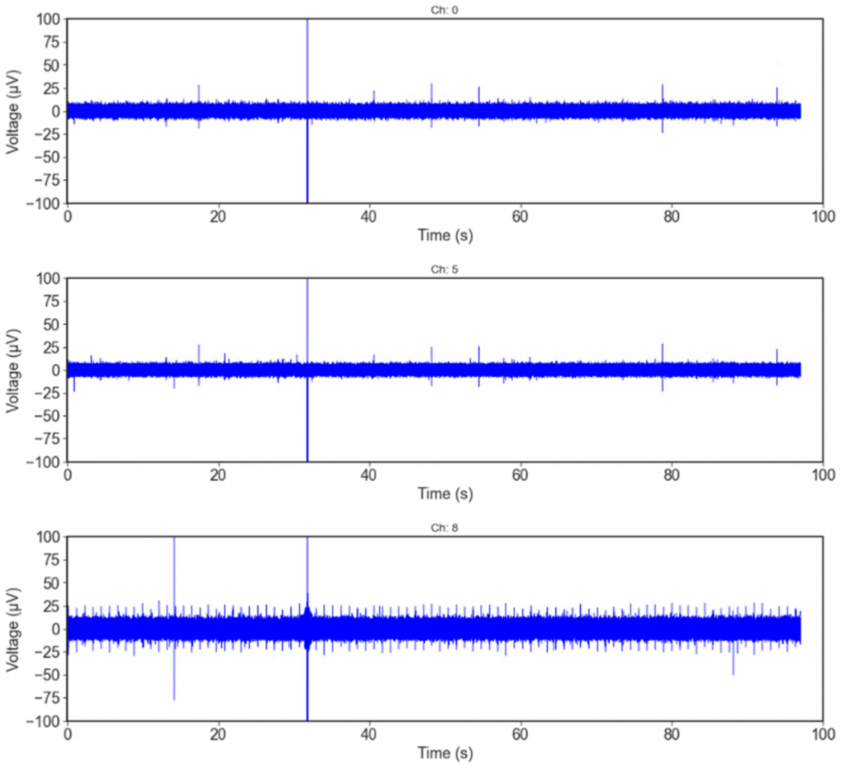


**Figure S11-2:** Subject 2 cVN spontaneous activity recording.

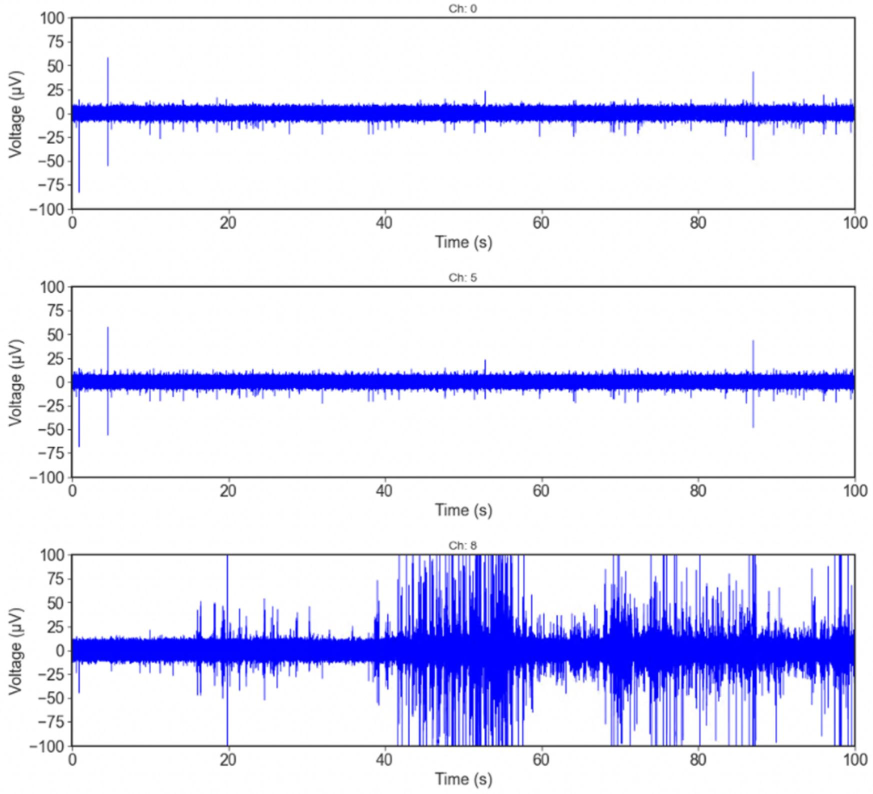


**Figure S11-3:** Subject 3 cVN spontaneous activity recording.

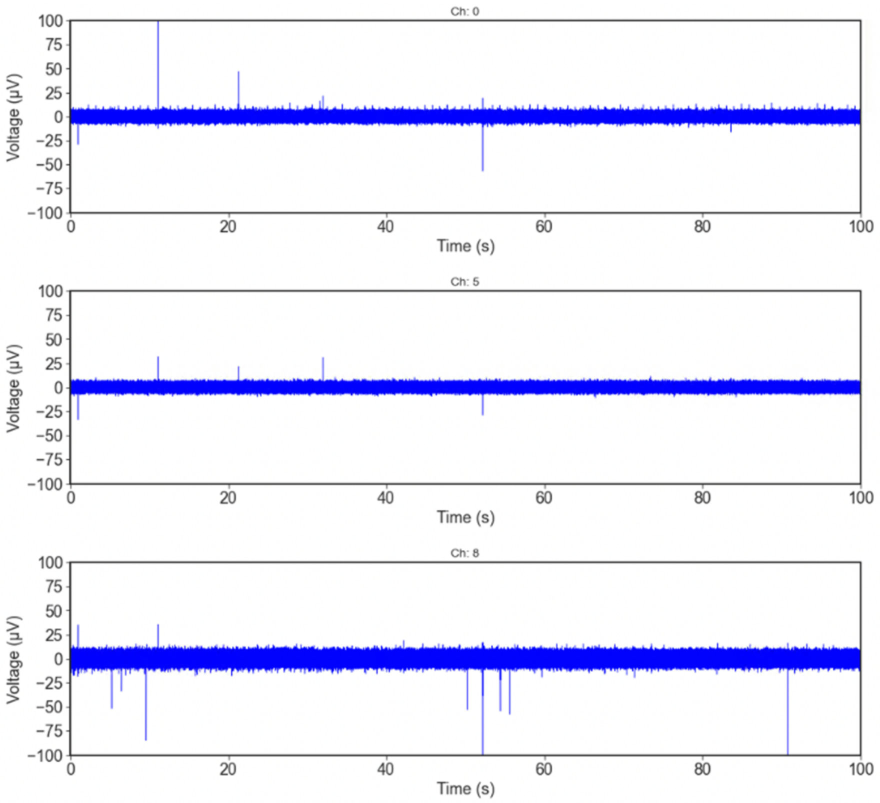


**Figure S11-4:** Subject 4 cVN spontaneous activity recording.

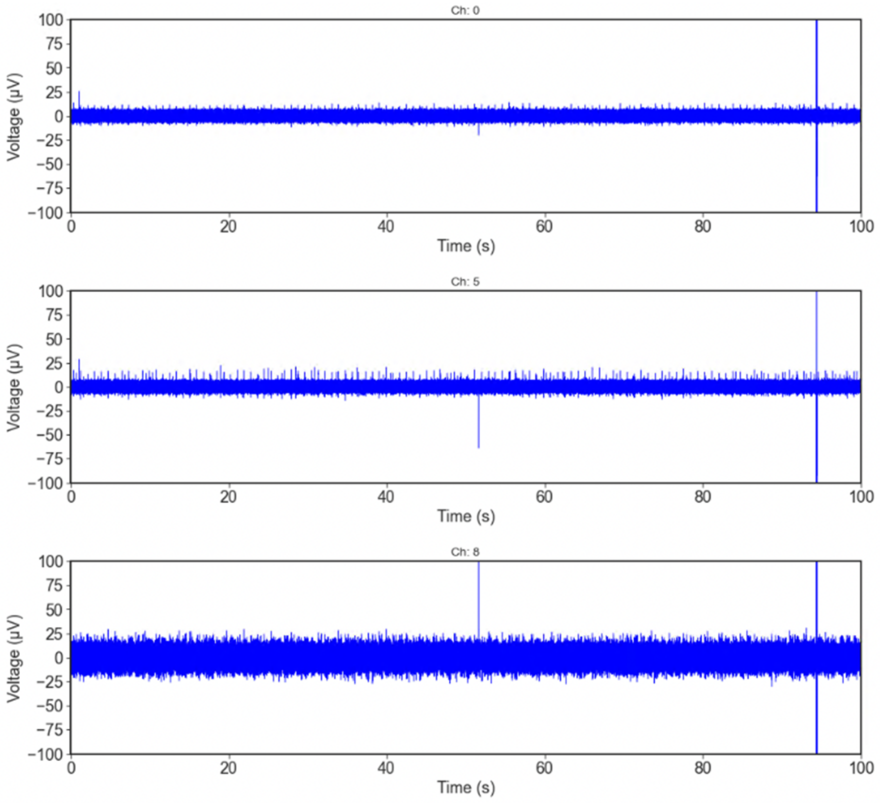


**Figure S11-5:** Subject 5 cVN spontaneous activity recording.

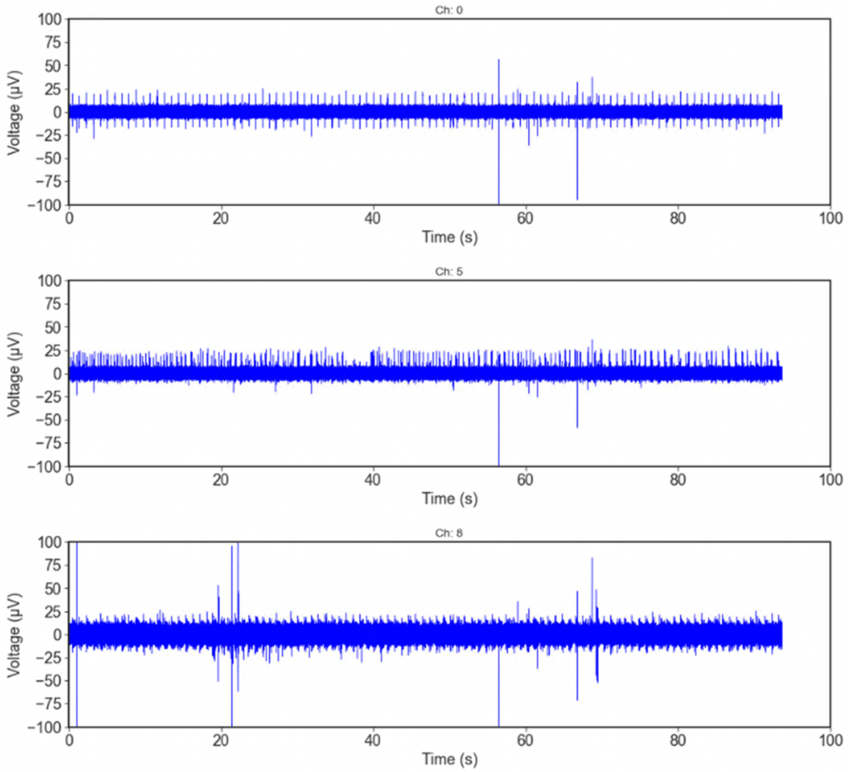


**Figure S11-6:** Subject 6 cVN spontaneous activity recording.
